# Complete telomere-to-telomere genomes of cowpea reveal insights into centromere evolution in Phaseoleae

**DOI:** 10.1101/2025.09.04.674167

**Authors:** Chuanzheng Wei, Shichao Sun, Yinzi Wang, Li Liu, Sofie Pearson, Yanbo Wang, Tashi Dorjee, Emma Mace, David Jordan, Yan Yang, Yongfu Tao

## Abstract

Cowpea (*Vigna unguiculata*) is a versatile legume crop providing a critical source of grain, vegetable and forage globally. Cultivated cowpea is classified into two main subspecies, subsp. *sesquipedalis* for fresh-pod vegetable and subsp. *unguiculata* for grain production. Here, we present two complete telomere-to-telomere (T2T) assemblies for the grain-type inbred lines HJD and vegetable-type FC6 through integrating PacBio HiFi reads, Oxford Nanopore ultra-long reads, and Hi-C data. The T2T genomes demonstrated improved contiguity, completeness, and accuracy compared to existing genomes, revealing clear telomeric and centromeric features. Comparative analysis of the T2T genomes highlighted inversions underlying subspecies divergence in cowpea. Evolutionary analysis uncovered contraction of gene families related to symbiosis in HJD, consist with its reduced root nodules compared to FC6. Distribution and composition of tandem repeat arrays and transposable elements in centromeric regions were largely conserved in cowpea, but displayed pronounced variation among Phaseoleae. Furthermore, frequent shifts of centromeric locations coincided with inversions found in Phaseoleae. Overall, this study provides a set of fundamental resources for cowpea improvement and enhances our understanding of cowpea subspecies divergence and genome evolution in Phaseoleae.

## Introduction

Cowpea (*Vigna unguiculata*) is an annual herbaceous species in the Fabaceae family, native to the semi-arid tropics of Africa [1,2]. It is widely cultivated across semi-arid tropical regions of Africa, Asia, and Latin America, due to its well-known adaptation to infertile soils and low-rainfall conditions [3,4]. Cultivated cowpea has two major subspecies, *unguiculata* and *sesquipedalis* [5]. The *unguiculata* subspecies is the second most grown legume in Africa for seed production, providing a critical source of protein for the most vulnerable people threatened by malnutrition [6,7]. The *sesquipedalis* subspecies, also known as the yardlong bean, is predominantly cultivated in East Asia for fresh pod consumption [8,9]. In addition, cowpea is also a popular forage crop due to its high protein content, digestibility and fast-growing character [10]. Therefore, genetic improvement of cowpea is critical for global food security.

Obtaining high-quality reference genomes is critical for exploring genomic variation to improve crop productivity [11]. The first reference genome of cowpea, IT97K-499-35, was assembled using Pacific Biosciences (PacBio) continuous long reads, covering 519 Mb DNA sequence [12]. Despite recent improvements, this reference genome is still highly fragmented with 46.0 Mb unanchored sequence and dozens of unfilled gaps [13]. Recent genome sequencing efforts have resulted in chromosome-scale assemblies of additional cowpea genomes [14,15]. However, these incomplete genomes still hinder a holistic view of genomic variation and further functional investigations. Complete genomes also present a unique opportunity to systematically characterize highly repetitive regions, such as centromeres and telomeres.

The Phaseoleae tribe, including numerous widely grown grain and forage species, represents an important lineage in the Papilionoideae subfamily of Fabaceae [16]. Phaseoleae is dominated by three major genera: *Phaseolus*, *Vigna*, and *Glycine* [17]. These closely related genera share strong macro–synteny with large conserved genomic blocks identified among their genomes [13,18–20]. Centromeres are crucial chromosomal functional domains that guarantee the faithful replication and precise separation of chromosomes during mitosis and meiosis [21]. Centromere repositioning events have been reported among *Vigna angularis*, *Vigna unguiculata*, and *Phaseolus vulgaris* through sequence comparison and oligonucleotide–FISH chromosomal painting [22,23]. However, centromere evolution and variation in Phaseoleae are still largely elusive due to the lack of high-quality references covering the highly repetitive regions. Nevertheless, the recent completion of telomere-to-telomere (T2T) genome assemblies for major Phaseoleae species has laid a solid foundation for in-depth investigation of centromere evolution [24–27].

In this study, we aim to 1) employ a combination of PacBio HiFi, Oxford Nanopore sequencing technologies to assemble complete T2T genomes of cultivated cowpea; 2) use multiple measures to demonstrate the superior quality of our T2T genomes and their potential in enhancing genetic discovery; 3) evaluate conservation and variation of repetitive elements and centromeres in Phaseoleae using the T2T genomes. This study will provide a set of fundamental genomic resources for cowpea improvement and shed new lights on centromere evolution in Phaseoleae.

## Results

### Telomere-to-telomere genome assembly and annotation of two cowpea lines

To assemble the genomes of HJD and FC6, we generated 37.33 Gb (∼73× coverage) of PacBio HiFi reads, 61.01 Gb (∼119× coverage) of Oxford Nanopore ultra-long reads and 32.12 Gb Hi-C reads for HJD. Additionally, we retrieved 28.59 Gb (∼57× coverage) of HiFi reads, 67.27 Gb (∼133× coverage) of ONT reads and 24.10 Gb Hi-C reads of FC6 from a previous study [15] (Fig. 1A, Table S1). K-mer analysis (k=21) based on HiFi reads estimated the genome size of HJD at approximately 481.93 Mb with a heterozygosity rate of 0.06%, while the FC6 genome was similarly sized at 474.17 Mb with a higher heterozygosity rate of 0.18% (Fig. S1, Table S2).

**Figure 1.**
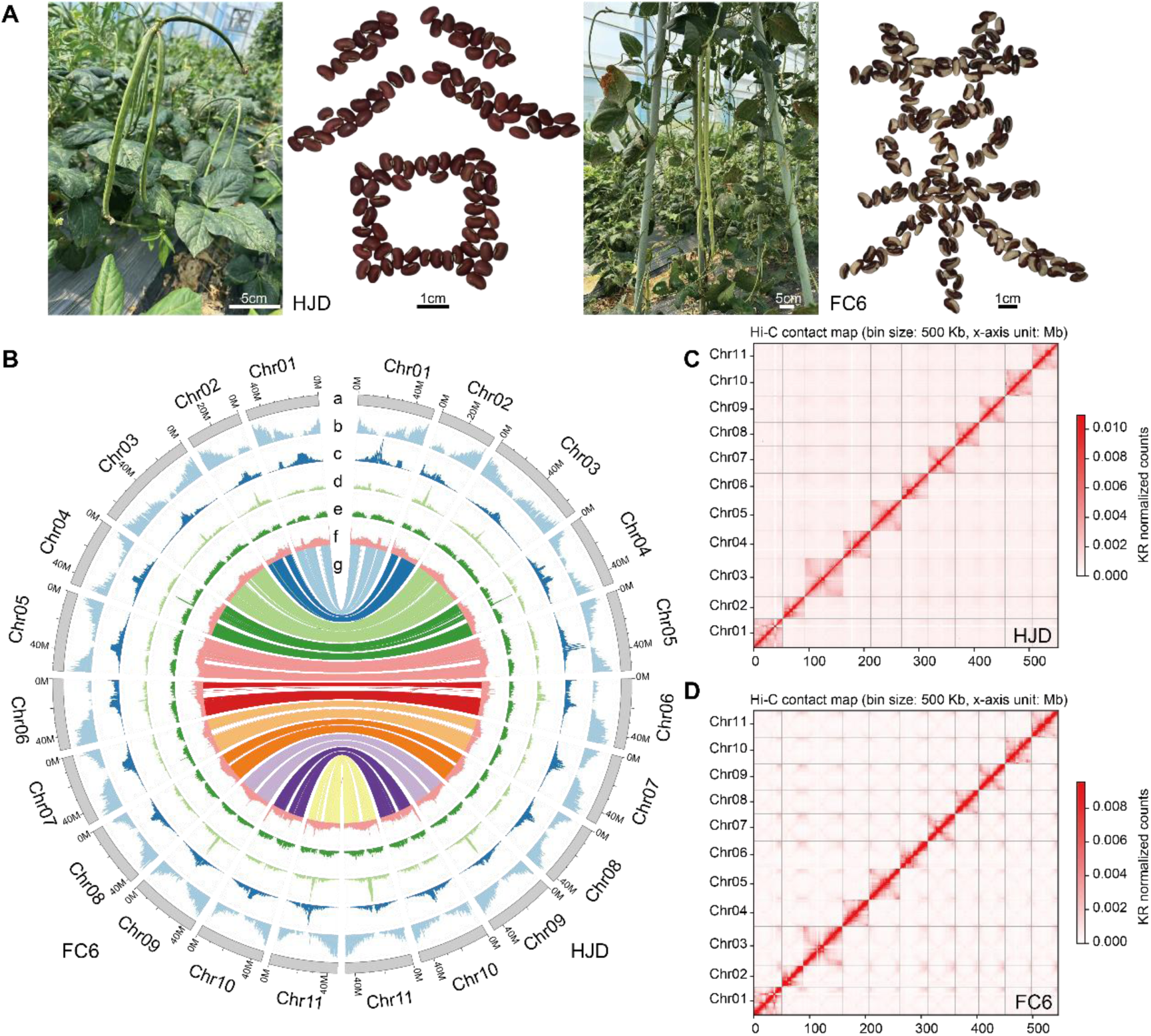
Assembly and Genomic Features of Grain-Type Cowpea HJD and Vegetable-Type Cowpea FC6. (A) Field-grown pods and seeds of HJD and FC6. (B) Circos plots of the HJD and FC6 assemblies showing: (a) chromosome layout; (b) gene density; (c) LTR/Gypsy element density; (d) LTR/Copia element density; (e) DNA transposon density; (f) GC content; (g) syntenic relationships among homologous genes. All tracks were computed using 500-kb windows. (C) Hi-C interaction maps at 500-kb bin size. Interaction frequencies are indicated by a color gradient from the plot periphery toward the diagonal, reflecting low to high contact intensity.

The preliminary assemblies were generated using Hifiasm by integrating PacBio HiFi, Oxford Nanopore ultra-long, and Hi-C reads. The draft assembly of HJD was 525.19 Mb in 15 contigs, and the FC6 genome amounted to 517.59 Mb in 17 contigs (Table S3). After removing organellar sequences, we employed HapHiC software’s quickview mode to orient and cluster contigs before manually refining the assignments in Juicerbox to anchor the assemblies onto 11 chromosomes. The resultant HJD assembly contained 4 unresolved gaps and 6 unassembled telomeric regions, while the FC6 assembly had 6 gaps and 3 missing telomeric regions. To fill the scaffold gaps, we used preliminary contigs generated by verkko and NextDenovo to close 2 gaps in HJD and 3 in FC6, and then closed remaining gaps using ONT ultra-long reads (Table S4). We appended all unassembled telomeric sequences following Liu et al. [28], yielding two gapless genome assemblies (Fig. 1B). The final HJD assembly reached 523.32 Mb with a contig N50 of 46.48 Mb, while FC6 totaled 517.39 Mb with a contig N50 of 46.29 Mb (Table 1).

**Table 1.** Cowpea genome assembly statistics.

|  | HJD | FC6 | IT97K-499-35 |
| --- | --- | --- | --- |
| Size of assembly (bp) | 548,737,058 | 542,521,633 | 519,435,864 |
| Contig N50 (bp) | 487,386,31 | 48,539,341 | 10,911,736 |
| Number of contigs | 11 | 11 | 754 |
| Scaffold N50 (bp) | 487,386,31 | 48,539,341 | 416,841,85 |
| Number of scaffolds | 11 | 11 | 686 |
| Number of Telomeres | 22 | 22 | 5 |
| Number of gaps | 0 | 0 | 68 |
| BUSCO of assembled genome (%) | 98.4 | 98.5 | 98.3 |
| Protein-coding genes | 27357 | 27670 | 31948 |
| BUSCO of protein-coding genes (%) | 98.4 | 98.4 | 98.6 |
| Repeat ratio in genome (%) | 51.80 | 51.26 | 48.35 |

Multiple strategies were employed to assess the accuracy and completeness of the T2T assemblies. Chromosome-level Hi-C heatmaps revealed high interaction intensity within each chromosome and no anomalous contact patterns, validating the proper ordering and orientation of all pseudomolecules (Fig. 1C-D). HiFi and ONT reads were remapped to the final assemblies, yielding mapping rates above 99.50% for all datasets except for ONT reads in FC6 with a mapping rate of 98.94% (Table S5). The uniform genome-wide coverage across all types of reads also supported the high-quality of our assembly (Fig. S2). Assembly completeness of the two genomes was assessed using Benchmarking Universal Single-Copy Orthologs (BUSCO), yielding scores of 98.40% for HJD and 98.50% for FC6 (Fig. S3). The LTR Assembly Index was used to evaluate completeness of long terminal repeat (LTR) sequences, scoring 13.56 and 13.55 for HJD and FC6, respectively. Using the k-mer-based method implemented in Merqury, quality values of >50.55 were calculated for both assemblies, corresponding to base accuracy of > 99.99% (Fig. S4, Table S6). Overall, these assessments demonstrated the excellent quality of our T2T assemblies.

After repeat masking, we predicted 27,357 genes in HJD and 27,670 genes in FC6 combining *ab initio* gene prediction with homology and RNA-seq evidence. A total of 23,176 allelic gene pairs were found between the two genomes, representing approximately 84% of both gene sets. Predicted proteins were queried against various databases, identifying matches for 26,984 HJD genes (98.64%) and 27,290 FC6 genes (98.63%) (Table S7).

### T2T genomes improve genetic analysis in cowpea

The complete sequence of our T2T genomes allows identification of telomeric and centromeric regions in cowpea. We found a total of 22 telomeric regions on the 11 chromosomes in each genome, spanning 535–24,326 bp in length (Table S8). In plants, centromeric regions are generally composed of arrays of tandemly repeated monomers [29]. We searched for the cowpea-specific satellite repeat CEN455 to localize centromeres in our T2T assemblies. Intact CEN455 arrays were identified on seven of the 11 chromosomes, while partial CEN455 arrays were found on the remaining chromosomes (Table S9). We analyzed the density of tandem repeat using quarTeT and manually merged the most repeat-dense regions with CEN455 hits to define centromere boundaries. The size of centromere varied markedly among different chromosomes in both genomes (Table S10). However, these centromeric regions are highly conserved between HJD and FC6.

Compared to the reference genome IT97K-499-35, our two T2T assemblies anchored approximately 50Mb of new sequence into chromosomes with improved contiguity, completeness, and accuracy (Fig. 2A-2B). Comparative synteny assessment of the T2T assemblies against IT97K-499-35 uncovered significant sequence omissions in the latter, particularly around centromeric regions (Fig 2C). Further comparisons of IT97K-499-35 with the T2T assemblies identified large inversions on chromosomes 3, 4, 5, 7, and 10 in IT97K-499-35, likely due to previous assembly errors (Table S11).

**Figure 2.**
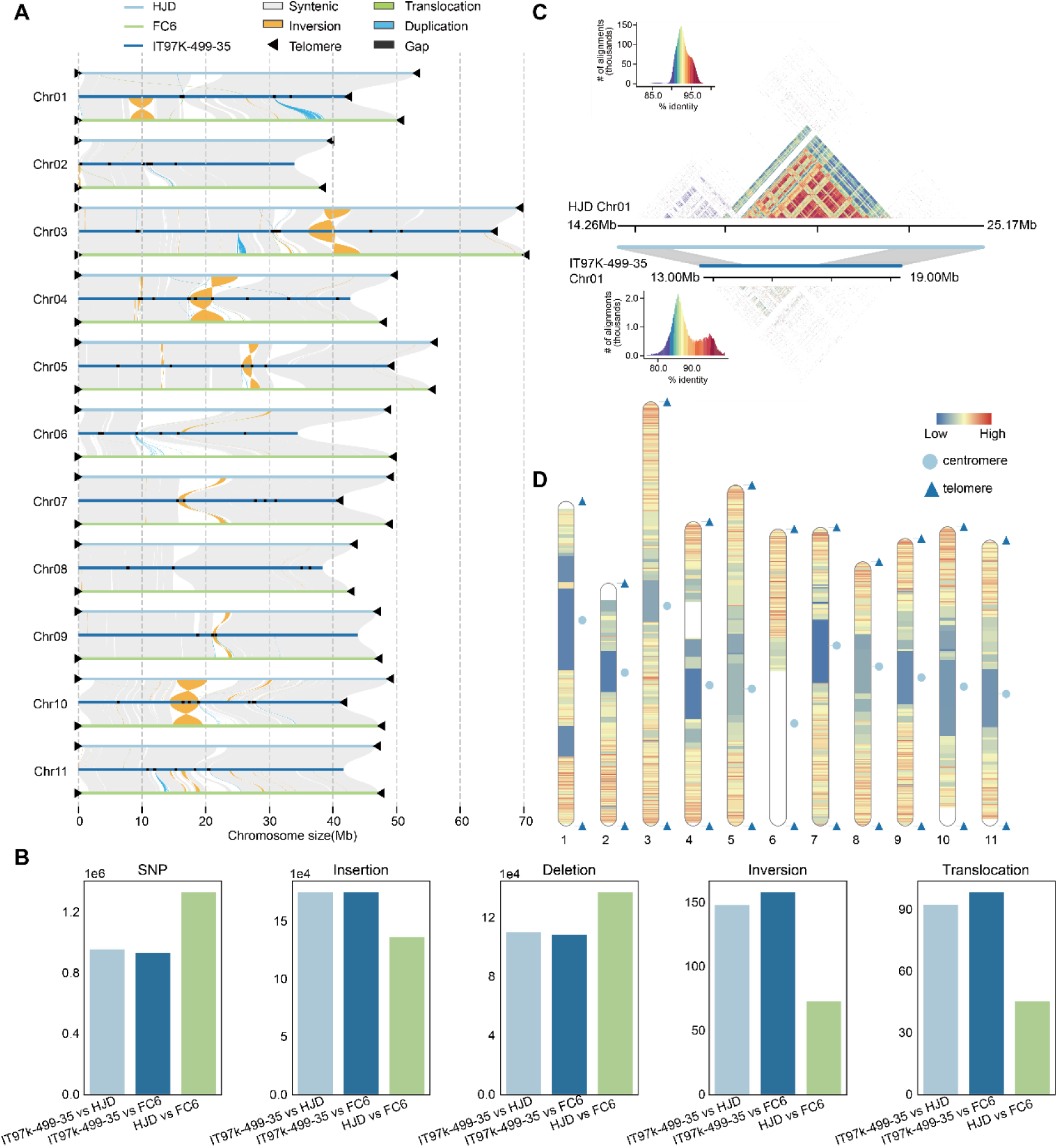
Comparison of cowpea T2T assemblies with IT97K-499-35 reference. (A) Syntenic overview of HJD (light blue), FC6 (dark blue) and IT97K-499-35 (cyan). Gray regions denote synteny, orange indicates inversions, green indicates translocations, teal indicates duplications, black indicates gaps, and black triangles mark telomeres. (B) Bar charts showing counts of SNPs, insertions, deletions, inversions and translocations for each genome pair (IT97K-499-35 vs. HJD, IT97K-499-35 vs. FC6, HJD vs. FC6). (C) Assembly improvement at the Chr01 centromere in HJD relative to IT97K-499-35. Gray blocks show synteny flanking the centromere; colored contact map displays similarity of tandem arrays. (D) Genome-wide recombination rate landscape across each chromosome of the HJD T2T assembly. Rates were calculated as genetic distance (cM) per physical distance (Mb) using linkage-map markers.

To assess the potential applications of our T2T genomes in genetic analysis, we firstly aligned a recently published cowpea genetic linkage map [12] to the HJD T2T assembly. Obvious marker intervals around centromeric regions were found on nearly every chromosome (Fig. 2D, Fig. S5-S6). The high-precision physical coordinates of our T2T assembly enabled accurate conversion of genetic distances to physical distances, and therefore, a refined assessment of recombination rate distribution across the genome. Secondly, re-sequencing reads of 270 cowpea accessions (45 *V. unguiculata* and 225 *V. sesquipedalis*) from a previous study were mapped to the T2T genomes to assess improvement in short-reads mapping [30]. The T2T assemblies reduced the number of unmapped reads by 33% compared to IT97K-499-35, with higher proportion of the genomes covered by these short-reads (Fig. S7, Table S12-S13). The mismatch rate also declined in the T2T genomes (Table S14). Thirdly, we identified 201 and 140 previously unannotated genes in the centromeric regions of HJD and FC6, respectively. GO functional annotation revealed that these genes were significantly enriched in multiple biological processes, including cellular respiration, immune response-activating cell surface receptor signaling, and positive regulation of defense response to insects (Fig. S8, Table S15), suggesting that centromeric regions contain genes critical for cowpea improvement.

### T2T genomes identified inversions influencing subspecies divergence in cowpea

Chromosomal inversions can play a critical role in racial divergence by markedly suppressing recombination and reducing genetic exchange [31]. Comparing the two cowpea T2T genomes identified 54 inversion events (Fig. S9). Nearly half (24/54) of these inversions showed pairwise fixation statistic (*F*_ST_) values between the cowpea subspecies higher than the genome wide average, suggesting their potential roles in driving subspecies differentiation (Table S16). In particular, the largest inversion spanning 9–13 Mb on Chr01 was unequivocally supported by Hi-C data (Fig. 3A). *F*_ST_ within this inversion reached 0.74, significantly higher than surrounding regions (Fig. 3B). Within each subspecies, nucleotide diversity (π) values inside the inversion were comparable to adjacent regions (Fig. 3C-D), indicating that the inversion region remains polymorphic. PCA based on SNPs within the inversion showed the two subspecies were clearly divided along the first principal component, which explained ∼89% of the variance (Fig. 3E). This evidence suggests that this inversion might play a critical role in the divergence of the two cowpea subspecies. GO enrichment analysis of genes within the inversion revealed significant overrepresentation of genes related to gibberellin biosynthesis and toxin metabolism pathways (Table S17).

**Figure 3.**
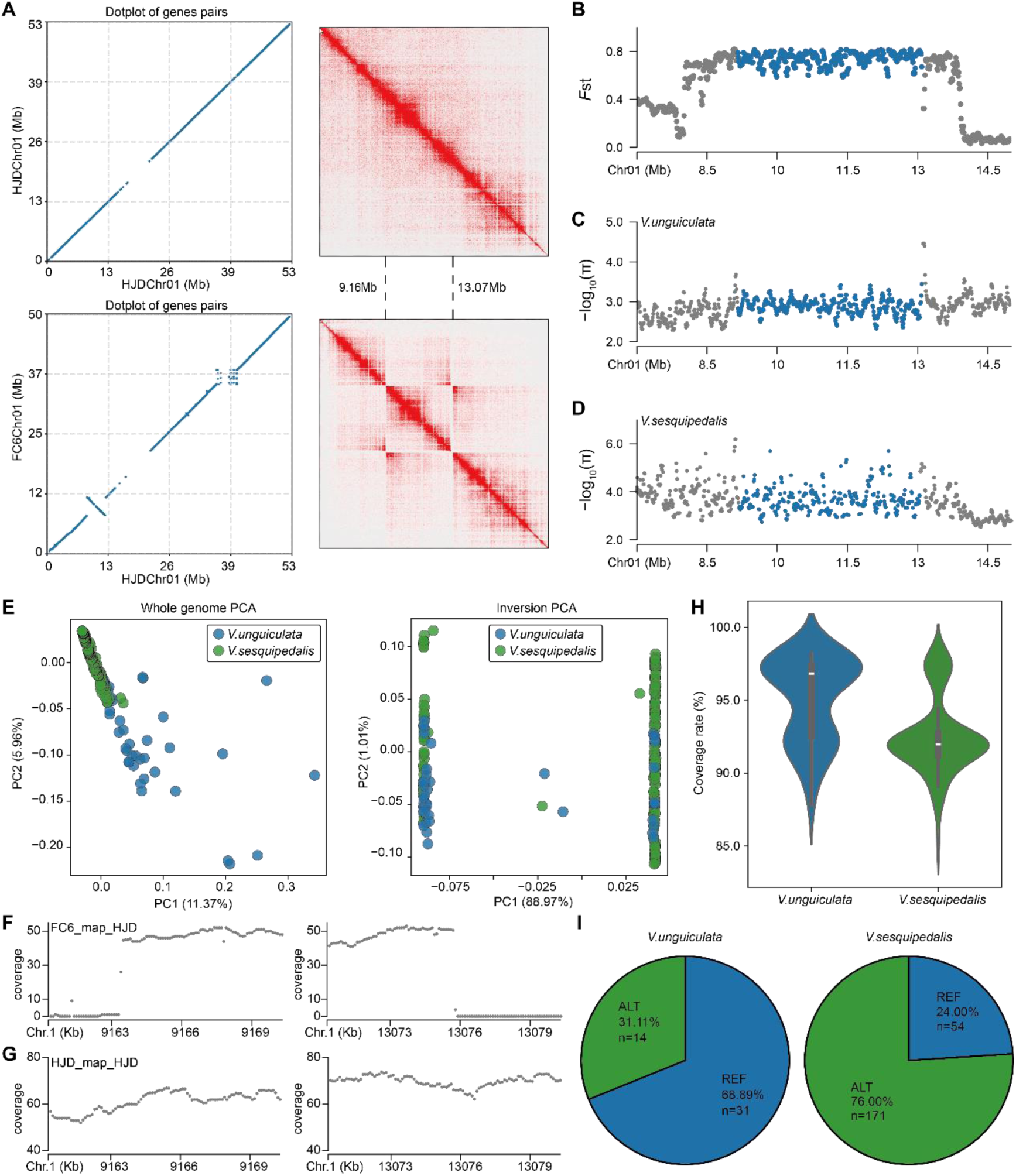
Inversion landscape on Chr01 and population genotyping. (A) Dot plot of syntenic gene pairs on Chr01 between HJD and FC6, alongside Hi-C contact matrices for the inverted region. Hi-C data from FC6 mapped to the HJD assembly reveal a butterfly pattern around the inversion. (B) Sliding-window F_ST_ distribution across the inversion and flanking regions, with the inversion highlighted in blue. (C) π distribution within the inversion and adjacent regions in *V. unguiculata*, inversion marked in blue. (D) π distribution for the same regions in *V. sesquipedalis*, inversion marked in blue. (E) Principal component analysis of genome-wide SNPs versus inversion-region SNPs. (F) Alignment gaps generated when mapping FC6 long reads to the HJD reference. (G) Continuous coverage profile of HJD long reads mapped to the HJD assembly. (H) Short-read coverage of population samples over the inversion region, using HJD as reference. (I) Distribution of reference and inverted genotypes within the population.

We further genotyped the inversion in the 270 cowpea accessions [30]. In the inversion region, *V. sesquipedalis* accessions showed pronounced coverage voids and reduced overall read depth (Fig. 3F), while *V. unguiculata* accessions retained consistently high coverage (Fig. 3G). Based on the different coverage patterns, the inversion was genotyped across the cowpea population (Fig. 3H). We found the HJD allele accounted for approximately 69% of *V. unguiculata* accessions compared to 24% of *V. sesquipedalis* (Fig. 3I). This marked divergence of allele frequency further supported the role of the inversion in subspecies differentiation in cowpea.

### Cowpea T2T genomes facilitated evolutionary analysis

Using the T2T genomes, we inferred the phylogenetic placement and divergence times of cowpea alongside 20 other plant species (Table S18). As expected, the phylogeny constructed from single-copy gene families was congruent with previously published trees. Molecular dating indicated that HJD and FC6 represented the most recently diverged lineages relative to IT97K-499-35. The split of cowpea from *Glycine* was dated at ∼22.62 Mya (Fig. 4A, Fig. S10), preceding soybean’s most recent whole-genome duplication (∼13 Mya) [32,33].

**Figure 4.**
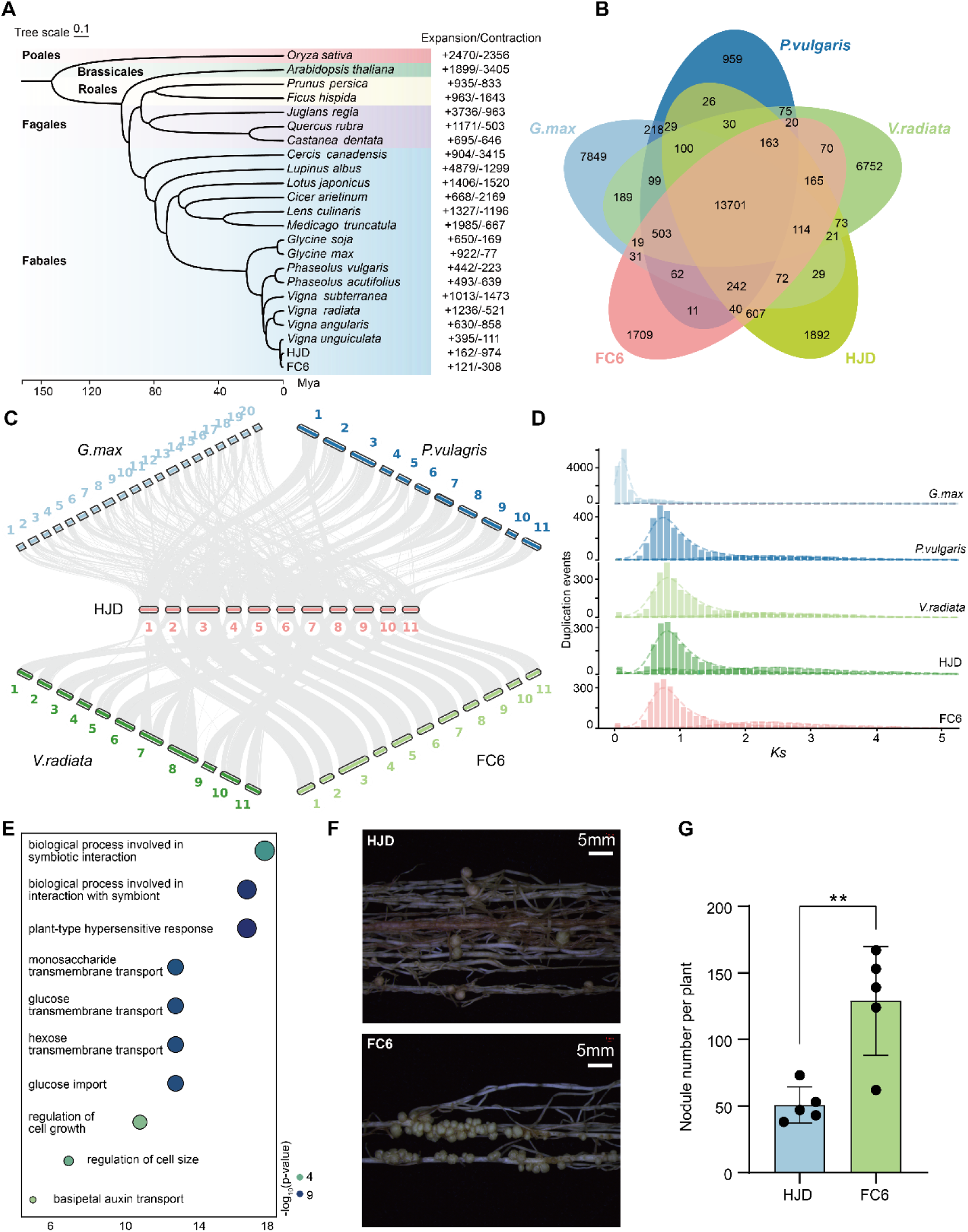
Comparative genomics and evolutionary analyses of cowpea. (A) Phylogenetic tree reconstructed from single-copy orthologs. Colored blocks (top to bottom) indicate Poales, Brassicales, Rosales, Fagales and Fabales. The timescale at the base shows divergence times, and numbers on the right indicate counts of contracted and expanded gene families. (B) Numbers of orthologous gene groups among Phaseoleae species (*G. max, P. vulgaris*, *V. radiata* and *V. unguiculata*). (C) Intergenomic syntenic blocks among Phaseoleae species. (D) Synonymous substitution (*Ks*) distributions for paralogous gene pairs in each species, where the y-axis reflects anchor-pair counts per Ks bin. (E) GO enrichment of contracted gene families in HJD. (F) Nodule phenotype of HJD 30 days after rhizobial inoculation. (E) Nodule phenotype of FC6 30 days after rhizobial inoculation. (G) Nodule counts for HJD and FC6 at 30 days (mean ± s.d., n = 5). Statistical significance between two groups was assessed by two-sided Student’s t-test; ** indicates *p* < 0.01.

A comparative analysis of four Phaseoleae species revealed 13,701 orthologous gene families conserved in all species and 3,601 gene families specific to cowpea (Fig. 4B). Synteny analysis among Phaseoleae species revealed that regions present as a single block in cowpea generally corresponded to two blocks in soybean, but only one block in mung bean and common bean (Fig. 4C). Within cowpea, a total of 194 syntenic blocks comprising 8,618 homologous genes (29.12% of the total) were identified, reflecting ancient polyploidy (Fig. S11, Table S19). Legumes trace their origin to a shared tetraploid ancestor, which underwent two polyploidization events at approximately 130 Mya and 58 Mya, with soybean subsequently experiencing an additional tetraploidization around 13 Mya [32,34]. The *Ks* distribution of soybean paralogs exhibited peaks at 0.59 and 0.12, corresponding to whole-genome duplication events dated to ∼58 Mya and ∼13 Mya. The only *Ks* peak in cowpea was at around 0.7, indicated no subsequent WGD events since ∼58 Mya (Fig. 4D, Fig. S10).

Further gene family evolution analysis identified expansions of 162 and 121 families in HJD and FC6, respectively, while 974 and 308 families exhibited contractions (Table S20). The expanded gene families in HJD and FC6 were enriched for different functions, potentially related to their distinct characters (Fig. S12). Intriguingly, GO terms associated with symbiosis and immune response were overrepresented in contracted gene families in HJD (Fig. 4E, Table S20). Our rhizobial inoculation assays revealed that HJD formed only ∼50% of the root nodules produced by FC6 (Fig. 4F-G). The reduced root nodules in HJD could be related to the contraction of genes related to symbiosis and immune response.

### Tandem repeat arrays are highly diverse among Phaseoleae species

Tandem repeat arrays (TRAs) are highly repetitive elements that are difficult to resolve in genome assemblies. The cowpea T2T genomes offered an opportunity to compare the distribution and composition of TRAs across Phaseoleae (Table S21). We found almost all TRAs overlapped with centromeric regions in *Glycine max* (Fig. S13). This concentration of TRAs around the centromeres was not observed in other Phaseoleae. In particular, eight out of 11 TRAs in *P. vulgaris* resided in non-centromeric regions. These TRAs exhibited extensive interspecific length variation, ranging from 12.89 Mb in HJD to just 0.59 Mb in *P. vulgaris* (Fig. S14-S17). The major component of TRAs, monomeric repeat units, showed distinct length-distribution profiles across the examined species. *G. max* had the shortest TRA monomers, predominantly clustering around ∼100 bp, followed by *V. radiata* and cowpea peaking at around ∼175 bp and 455bp, respectively (Fig. 5A and 5B). *P. vulgaris* showed three peaks of monomer lengths at ∼100 bp, ∼270 bp, and ∼530 bp (Fig. 5A).

**Figure 5.**
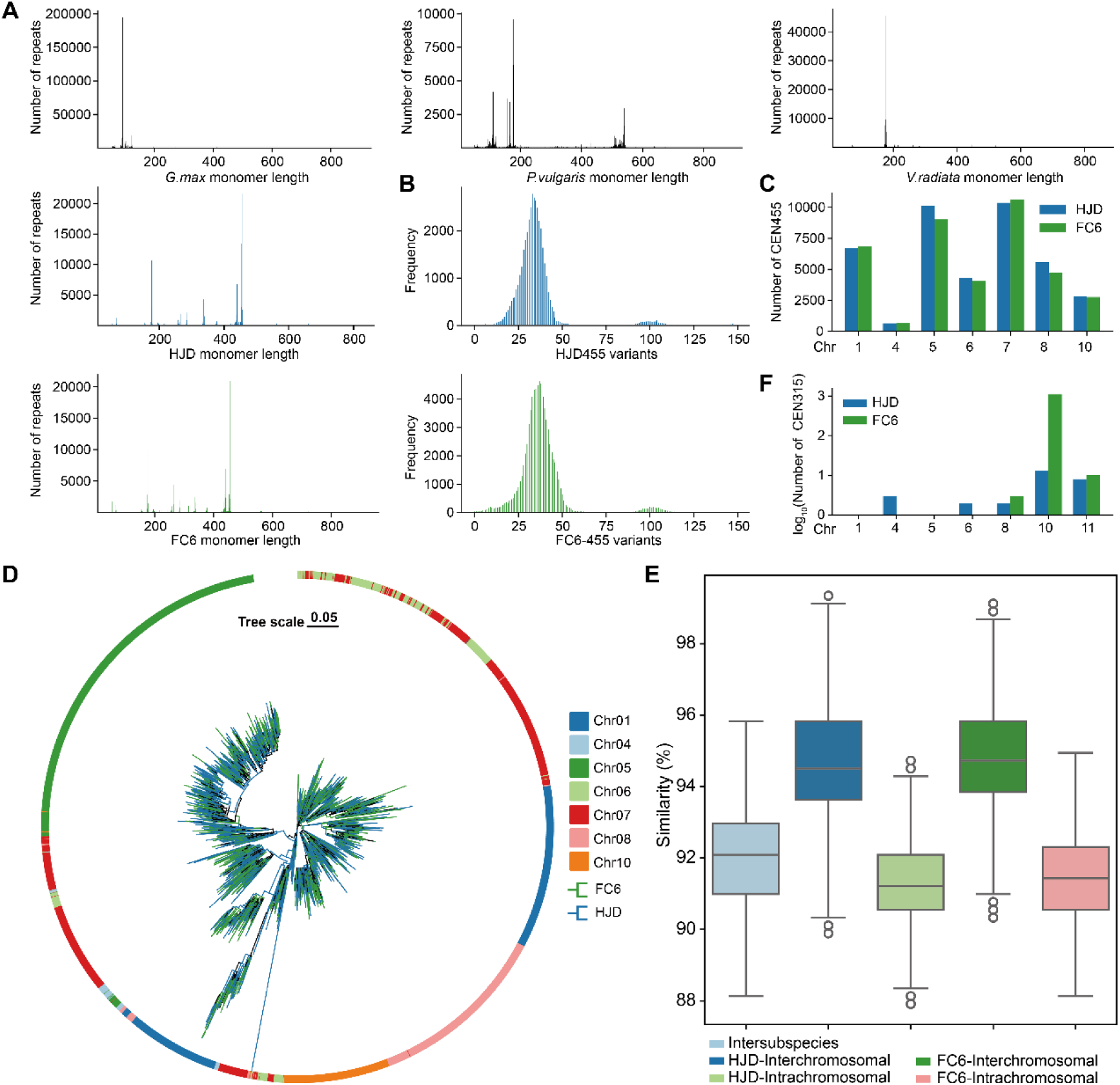
Characteristics of Phaseoleae TRAs. (A) Length distribution of monomeric tandem repeats identified by TRASH in each genome. (B) Distribution of variation counts in CEN455 compared with the consensus monomer. (C) Copy number of CEN455 on each chromosome in HJD and FC6. (D) Copy number of CEN315 on each chromosome in HJD and FC6. (E) Phylogenetic tree of sampled CEN455 monomers. Outer ring colors denote chromosomes; inner branch colors distinguish HJD and FC6. (F) Boxplot comparison of CEN455 monomer similarity across subspecies, within chromosomes, and among chromosomes.

CEN455 is the dominant monomeric repeat units in cowpea. A total of 40,566 and 38,720 CEN455 repeat units were identified in HJD and FC6, respectively (Fig. 5C, Fig. S13). An evolutionary tree showed that CEN455 repeats segregate mainly according to their chromosomal provenance rather than subspecies (Fig. 5D). CEN455 repeat units showed elevated sequence similarity within a genome (Fig. 5E). The satellite repeat CEN440 displayed a similar pattern to CEN455 (Fig. S18). In contrast, CEN315 repeats were detected almost exclusively in FC6 (Fig. 5F).

Sequence similarity was observed among different classes of TRAs within species, such as, Gm103 and Gm123 in *G. max*, Pv111 and Pv166 in *P. vulgaris*, and CEN440 and CEN455 in cowpea. Additionally, interspecies homology was detected among repeats including Gm335, Vr215, Pv539, and CEN338. Phylogenetic inference indicates these satellites might have derived from a shared ancestral repeat and undergone lineage-specific expansion following speciation (Fig. S19). Our results underscore substantial interspecific variability of TRAs within Phaseoleae.

### Comparison of transposable elements reveals Phaseoleae divergence

To investigate the impact of transposable element (TE) dynamics on genome evolution in Phaseoleae, we quantified TE content and composition. TEs comprised roughly 50% of each genome in all four species, despite their differences in genome size. LTR retrotransposons was the dominant component of TE, accounting for 31%–40% of the genome sequence (Fig. 6A, Table S22). TE accumulation accounted for nearly half of genome size variation in these species (Fig. 6B, Table S23). Interestingly, the proportion of specific TE families differed markedly among species with comparable TE content. Bianca-type Copia was nearly exclusively found in cowpea. Within Gypsy elements, CRM was the most abundant subfamily in cowpea, whereas Athila and Retand predominated in mung bean and common bean, respectively (Fig. 6C-D, Table S24).

**Figure 6.**
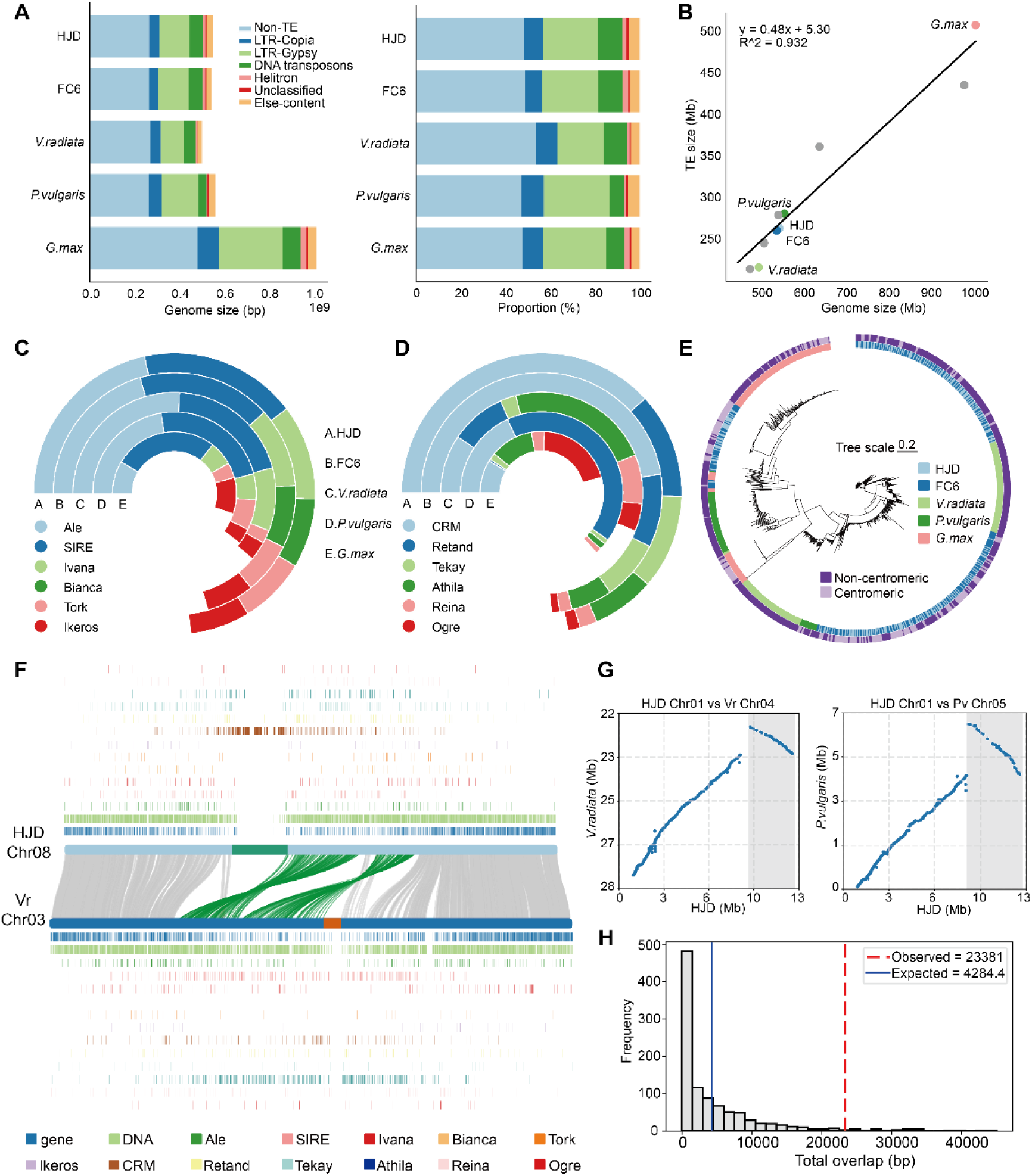
Analysis of repetitive sequences in Phaseoleae. (A) Genome composition of each species and proportions of repeat classes. (B) Relationship between transposable element content and genome size. (C) Distribution of Copia superfamily members. (D) Distribution of Gypsy superfamily members. (E) Phylogenetic tree of full-length CRM elements. (F) Synteny between cowpea and mung bean; central lines connect homologous chromosomes, colored blocks mark centromeres, gray lines link homologous genes, and green lines highlight genes within or adjacent to centromeres. (G) Dot plots of inversions at shared breakpoints in cowpea versus mung bean and cowpea versus common bean; blue dots represent orthologous gene pairs, gray shading indicates aligned regions. (H) SD occupancy at inversion breakpoints estimated from 1000 random resampling iterations.

We further examined centromeric sequence composition, which showed that TEs occupied 24.88%–98.78% of centromeric regions. Gypsy elements were the predominant TE class in these regions, accounting for 69.29%–88.38% of centromeric TEs. The CRM subfamily represented the major component in cowpea centromeres, comprising 61.42% and 52.53% of centromere sequence in HJD and FC6, respectively (Table S25). A phylogeny based on CRM elements showed that CRM monomers from different species cluster together within multiple strongly supported lineages, suggesting recurrent incorporation of CRM into centromeric arrays (Fig. 6E). In *V. unguiculata*, CRM repeats were tightly interspersed across both centromeric and non-centromeric clades, reflecting recent bursts of element amplification in cowpea. In contrast, soybean CRM elements formed well-resolved lineages, consistent with an ancient expansion event in *G. max* (Fig. 6E). These patterns indicate that CRM elements have retained ancestral diversity while undergoing lineage-specific expansions throughout Phaseoleae evolution.

### Frequent Centromere Renewal and Repositioning in Phaseoleae

Comparison of centromere positions in Phaseoleae found frequent centromere renewal and repositioning. For example, the centromere of cowpea Chr08 did not align with the syntenic mung bean Chr03 (Fig. 6F). Changes in arm ratio of corresponding chromosomes in Phaseoleae species further illustrated centromere repositioning. The arm ratios for the syntenic group consisting of cowpea Chr07, mung bean Chr10, and common bean Chr07 were 1.64, 2.01, and 2.57, respectively. Remarkably, centromere repositionings coincided with nearby inversion events in all 11 syntenic chromosome pairs between cowpea and mung bean and in 10 out of 11 pairs between cowpea and common bean (Fig. S20-S21). This suggests that centromere repositioning is likely driven by inversion events.

Multiple centromere inversions shared nearly identical breakpoints, such as these breakpoints on Chr01 (∼13 Mb), Chr03 (∼21 Mb), and Chr06 (∼1 Mb) of HJD (Fig. 6G, Fig. S22). Repetitive elements are a major force driving inversion formation. Therefore, we further assessed whether TEs and segmental duplications (SDs) were associated with inversion breakpoints (Table S26). SDs were found to be highly enriched at breakpoints of these inversions, indicating they play a role in inversion formation around centromere and centromere repositioning (Fig. 6H, Fig. S23).

## Discussion

High quality reference genomes are fundamental resources for genomic investigation and genetic improvement of plants. This study presented the first two complete T2T genome assemblies of cowpea, resolving previously intractable regions, including centromeres and telomeres. These T2T genomes showed much higher contiguity, completeness, and accuracy compared to previous cowpea genomes. The improved mapping statistics in analysis of re-sequencing data demonstrated that the superior quality of our T2T genomes can enhance genetic discovery. Comparative analysis of our T2T genomes found that inversions could play an important role in cowpea subspecies divergence. The holistic view of centromeres and repetitive elements from the T2T genomes uncovered frequent centromere renewal and repositioning in Phaseoleae, which often coincided with inversions. In summary, the findings of this study provided a set of foundational resources for cowpea improvement and enhanced our understanding of cowpea diversification and genome evolution in Phaseoleae.

T2T genomes provide new opportunities to generate highly accurate and complete references to explore the full genomic landscape in a species. Recent advances in sequencing technology has enabled assembly of T2T genomes for multiple species, including rice [35], maize [36], sorghum [37], soybean [24], common bean [26] and mung bean [27]. These genomes uncovered genomic sequence in highly complex and previously intractable genomic regions and have proved their value in facilitating investigation of genomic pattern and gene mining. Likewise, the T2T genomes in this study captured full genome sequence, including centromere regions in cowpea. The identification of hundreds of new protein-coding genes in centromeres of our cowpea T2T genomes with GO enrichment in immunity and stress responses demonstrated the functional importance of these regions. Access to the T2T genomes provides a solid foundation for full-scale exploration of genomic variation to improve cowpea productivity.

The T2T assemblies have enabled the detailed examination of centromeres in cowpea and comparison among Phaseoleae species. We found composition and position of centromeres are largely conserved within cowpea. In contrast, Phaseoleae species displayed frequent repositioning of centromere and distinct centromere composition. Rapid centromere evolution is well-documented in previous studies [38]. Despite their conserved function, centromeres show pronounced variation in terms of sequence components and physical position in *Gossypium hirsutum* [39] and *Fragaria x ananassa* [40]. Centromere-specific proliferation of TEs can create new binding sites for centromere proteins and disrupt old ones, thereby causing positional shifts of centromeres [41,42]. Our results supported the role of TEs as key drivers of sequence variation and movement of centromeres in Phaseoleae. Similarly, centromere relocation promoted by pervasive TE insertions was reported in Salicaceae species [43]. In addition, the enrichment of SDs at the inversion breakpoints around centromere suggests that duplication-mediated inversions could also play a role in centromere shifts. Overall, our examination of centromeres in Phaseoleae underscores that centromeres are hotspots of genomic innovation.

Comparison of our cowpea T2T assemblies inidcated that inversions likely underlie the divergence between the two cultivated subspecies. In particular, the ∼4 Mb inversion on chromosome 1 showed clear allele frequency difference between *V. unguiculata* and *V. sesquipedalis*. Chromosomal inversions can prevent gene exchange by suppressing recombination in the region, therefore locking together co-adapted allele combinations [31]. This mechanism is well-recognized as a driver of speciation and racial divergence. Large chromosomal inversions related to population divergence have been reported in *Sesbania bispinosa* [44] and Lake Malawi cichlid [45]. Nevertheless, it is still unclear what traits are affected by the large inversion in cowpea. GO enrichment analysis showed genes in the inversion region were primarily enriched for gibberellin biosynthesis and toxin metabolism (Table S17), which are likely related to the different plant architecture and growing environments of the two subspecies. Future functional investigation of these genes will clarify how this inversion contributed to cowpea diversification.

In summary, we generated complete T2T genome assemblies for the grain-type cowpea HJD and the vegetable-type cowpea FC6. The superior quality of the T2T genomes and their potential in improving genetic discovery were demonstrated. The T2T genomes enabled identification of inversions influencing subspecies divergence in cowpea and enhanced our understanding of evolutionary history of Phaseoleae. Analysis of repetitive sequence highlighted frequent centromeres changes with extensive content variation among Phaseoleae species. The T2T genomes in our study provide accurate and complete references for cowpea genetic improvement.

## Materials and methods

### Plant materials and sequencing

The cowpea cultivar HJD selected for genome sequencing was cultivated under greenhouse conditions for 30 days. Genomic DNA was extracted from young leaves using an improved CTAB method [46], and DNA integrity was verified by electrophoresis on a 0.75% agarose gel. High quality genomic DNA was then prepared with the Pacific Biosciences SMRTbell Express Template Prep Kit 2.0 following the manufacturer’s standard protocol. Each SMRTbell library was generated using the Pacific Biosciences SMRTbell Express Template Preparation Kit 2.0. The constructed libraries underwent size-selection for approximately 15 Kb fragments using BluePippinTM, and polymerase binding to the SMRTbell templates was performed using the DNA Polymerase Binding Kit. Subsequently, SMRT cell libraries were constructed and sequenced using the Pacific Biosciences Sequel II sequencing platform. For Oxford Nanopore ultra-long sequencing, high-molecular-weight genomic DNA fragments were enriched using the Short Read Eliminator XL following Oxford Nanopore’s ultra-long sequencing guidelines. Libraries were then constructed using the Oxford Nanopore SQK-LSK110 Kit as per the standard protocol and sequenced on the PromethION platform. Hi-C libraries were constructed following previously established procedures [47], and sequencing data were generated using the Illumina HiSeq X Ten platform.

### Genome assembly and quality evaluation

Genome sizes and heterozygosity for the HJD and FC6 genomes were estimated by Jellyfish v2.2.10 [48] in combination with GenomeScope2 [49]. Initial genome assemblies were generated using hifiasm v0.20.0 [50] in hybrid mode, integrating HiFi reads, ONT reads, and Hi-C sequencing data. Hi-C data facilitated scaffold anchoring and removal of low-quality short contigs. Following read mapping onto the initial assembly using BWA-mem v0.7.17 [51], scaffolds were quickly ordered and oriented using the quick view mode of HapHic v1.0.6 [52]. Further refinement of genome assemblies was performed using interaction signals visualized in Juicebox, generating scaffolds with identified gaps. Concurrently, two additional initial assemblies were independently generated using verkko v1.4.1 [53] and NextDenovo v2.5.2 [54], which were subsequently employed for genome gap filling. Genome gaps were further filled using Winnowmap v2.03 [55]. For each gap bridged perfectly by both HiFi and ONT long reads, the longest and highest-quality alignment was chosen to replace the missing sequence and close the gap. Missing telomeric sequences were supplemented according to the procedure described by Liu et al. [28]. Briefly, hifiasm was used to assemble two read sets separately: (1) reads not mapped to the genome and (2) HiFi reads containing telomeric repeats (TTAGGG). The assembled telomeric contigs were then aligned to the chromosome scaffolds with minimap2 v2.28, leveraging sequence overlaps, to generate a final genome assembly that incorporates complete telomeric sequences. After polishing with NextPolish v1.4.1 (https://github.com/Nextomics/NextPolish), the assembled HJD and FC6 genomes were compared against the previously published IT97K-499-35 reference genome to ascertain chromosome numbering and orientation. Ultimately, gap-free genome assemblies were achieved, and genomic interaction heatmaps were visualized using the plot mode of HapHic v1.0.6 [52].

Genome completeness was assessed using BUSCO v5.5.0 [56] based on the Fabales single-copy ortholog dataset (fabales_odb10). Genome assembly accuracy was evaluated by mapping whole-genome sequencing reads onto the HJD and FC6 assemblies using minimap2 v2.28 [57]. Alignment rates and coverage were calculated with pandepth v2.26 [58], and assembly quality values (QV) were estimated using the k-mer-based Merqury software [59]. Genome continuity was assessed by calculating contig N50 values, while genome assembly quality was further evaluated using the LTR Assembly Index (LAI).

### Repetitive element annotation

HiTE v3.2.0 [60] was utilized for high-accuracy identification of full-length TEs, employing a rapid and precise dynamic boundary adjustment strategy with parameters: --plant 1 --recover 1 --annotate 1. HiTE integrates the strengths of both *de novo* and signature-based approaches, fully exploiting TE repeat characteristics, conserved motifs, and structural features to achieve precise detection and generate a high-quality repeat library. TE classification at the superfamily level was performed using TEsorter v1.4.7 [61], with LTR-RTs further subclassified into distinct lineages. SDs in each assembled genome were identified using BISER v1.3 [62] with default parameters, employing the soft-masked versions of the respective genomes as inputs. We defined SD as repetitive segments longer than 1 kb exhibiting ≥90% sequence identity.

### Genome annotation

Following soft-masking of repetitive sequences, protein-coding genes were comprehensively predicted and annotated through an integrative approach, combining *ab initio* gene prediction, homology-based searches, and RNA-seq assembly-based predictions. Protein sequences from *Lotus japonicus*, *Medicago truncatula*, *Glycine max*, *Phaseolus vulgaris*, and *Vigna radiata* were employed as the protein family database. RNA-seq reads were mapped onto the reference genome using HISAT2 v2.2.1 [63] with default parameters. All data were integrated into the fully automated BRAKER3 [64] pipeline to train gene prediction models and accurately annotate genes via GeneMark-ETP and AUGUSTUS. Transcripts were assembled de novo using StringTie v2.2.1 [65], and the PASA v2.5.3 [66] software tool was utilized to predict and refine gene structures from these assembled transcripts through two iterative rounds, resulting in the final gene annotation models.

The completeness of the final protein dataset was assessed using BUSCO v5.0.0 in protein mode with the fabales_odb10 dataset. Functional annotation of protein-coding genes was initially conducted via Diamond Blastp (v2.1.9, --evalue 1e-5) [67] searches against UniProt, NR, GO, and KEGG databases. Additionally, conserved protein sequences, functional motifs, and structural domains were characterized by employing InterProScan and Hmmscan searches against the InterPro and Pfam databases.

### Identification of centromeres and telomeres

Telomeric regions were identified using seqtk v1.4(https://github.com/lh3/seqtk) by searching for canonical 5’-CCCTAAA and 3’-TTTAGGG repeats with the parameter seqtk telo -m CCCTAAA. Centromeric regions were defined based on the previously identified 455-bp centromere-specific satellite sequence in *V. unguiculata* by fluorescence in situ hybridization (FISH). Specifically, Blastn v2.14.0+ was used to search for these satellite repeats throughout the genome, and candidate regions were selected using a ≥90% sequence identity threshold. Given the observed high density of short tandem repeats and low gene density typical of centromeric regions, genome-wide sequence identity was computed using StainedGlass v0.6 [68]. Additionally, TRASH v1.2 [69] was used to identify and extract tandem repeats, with higher-order repeats recognized as pairs of highly similar repeat blocks. Approximate centromeric boundaries were determined based on consensus results obtained from multiple analytical methods. The centromere locations in soybean were sourced from previous research [24].

### Hi-C data analysis for inversion

Hi-C reads from HJD and FC6 were each aligned to the HJD reference genome using BWA v0.7.17 [51]. Optical duplicates, unmapped reads, and singleton mates were filtered out to produce the final alignment files with the command: bwa mem -5SP -t 48 HJD.fasta reads1 reads2 | samblaster | samtools view - -@ 48 -S -h -b -F 3340. Juicer was then used to generate .assembly and .hic files for visualization. Juicebox heatmaps of the inversion regions revealed the characteristic “butterfly” interaction pattern indicative of inversion events.

### Variant calling and population genetic analyses

A total of 270 genomic resequencing datasets (NCBI accession number PRJNA890023) were retrieved and employed for population genetic analyses. Raw sequencing reads were filtered with fastp v0.23.4 and aligned to the HJD genome assembly using BWA v0.7.17, followed by sorting and indexing of alignment files with SAMtools v1.21. SNP calling was performed using GATK v4.6.2 [70], and variants were filtered to retain high-confidence SNPs according to the criteria: QD < 2.0 || QUAL < 30.0 || SOR > 3.0 || FS > 60.0 || MQ < 40.0 || MQRankSum < -12.5 || ReadPosRankSum < -8.0. A total of 7,685,972 SNPs were retained for subsequent analyses. For principal component analysis (PCA), SNPs were further filtered using VCFtools v0.1.16 (MAF ≥ 0.03 and missing data rate ≤ 0.1), and PCA was conducted with PLINK v1.90b7 [71]. π and *F*_ST_ were calculated and analyzed using VCFTools v0.1.16 [71] with a window size of 50,000 bp.

### Genotyping of inversions within populations

HiFi sequencing reads from HJD and FC6 were independently aligned to the HJD genome using minimap2 v2.28. The genomic coverage within inverted regions was notably higher in the HJD_map_HJD combination compared to FC6_map_HJD. Alignment of Illumina resequencing reads revealed markedly distinct patterns of genomic coverage within inversion regions between *V. unguiculata* and *V. sesquipedalis*. Regions exhibiting genomic coverage >94% were classified as the reference genotype (no inversion), whereas lower coverage indicated the presence of inversions. PCA was performed using 19,754 SNPs from inversion regions, and inversion genotypes were assigned based on observed clusters. When discrepancies between the two genotyping approaches occurred, inversion genotypes were manually resolved using IGV v2.16.2 [72] by inspecting read coverage patterns near inversion breakpoints.

### Collinearity analysis of different genomes

HJD and FC6 genome assemblies were aligned against the reference genome IT97K-499-35 (retrieved from https://phytozome-next.jgi.doe.gov/) using minimap2 v2.28 [57]. Syntenic regions and structural variations between genomes were identified using Syri v1.6.5 [73]. Structural variant statistics (number and length) were calculated with custom scripts, and genome alignments were visualized using plotsr v1.1.1 [74].

### Comparative genomics analysis

Single-copy orthologous protein sequences were identified with OrthoFinder v2.5.5 [75], aligned independently with MUSCLE v5.1, and then concatenated to form a super-alignment matrix. A phylogenetic tree encompassing 21 species was constructed using RAxML with the maximum likelihood method, supported by 100 bootstrap replicates. Divergence time estimation based on the phylogenetic tree was performed using the MCMCTree program in PAML [76] with the following parameters: burn-in period = 10,000, number of samples = 100,000, and sampling frequency = 2. Furthermore, gene family expansion and contraction analyses were carried out using CAFÉ [77], and functional annotation of these gene families was conducted using clusterProfiler [78].

Gene-based synteny analyses were performed using JCVI v1.5.2 [79] to identify orthologous and paralogous genes among genomes. Synonymous substitution rates (*Ks*) of homologous gene pairs, commonly used to infer whole-genome duplication (WGD) events, were calculated and visualized using WGD v2.0.38 [80].

### Rhizobial inoculation experiment

Seeds of HJD and FC6 were surface-sterilized using chlorine gas for 12–14 h before being planted in pots and cultivated in a growth chamber under a 16-h light/8-h dark cycle at temperatures of 27℃ (day) and 22℃ (night). Seedlings were inoculated with *Bradyrhizobium japonicum* USDA110 at day 7 post-germination. *B. japonicum* was cultured in liquid YMA medium at 28℃ prior to inoculation. Each seedling was inoculated with 5 mL of bacterial suspension (OD₆₀₀ = 0.1 prepared in distilled water), and the number of root nodules was recorded after 30 days.

### Tandem repeat annotation and analyses

The TRASH pipeline v1.2 [69] was employed to identify tandem repeat arrays across all genomes. Genome-wide internal sequence similarities were computed and visualized using StainedGlass v0.6 [68] using default parameters. Monomer diversity was evaluated based on variant sites within repeat monomers. Using CEN455 as an example, all identified monomers were initially aligned using MUSCLE v5.1 to generate a consensus sequence, and a custom script was employed to determine the number of variant bases at corresponding positions relative to the consensus. Similarity was defined based on the ‘identity’ value obtained from Blastn alignments. To investigate the evolutionary origins of monomers, proportional random sampling by species and chromosome was performed to obtain approximately 1,800 monomers. Maximum-likelihood phylogenetic trees were subsequently constructed using FastTree v2.2 (https://github.com/morgannprice/fasttree), and tree structures were visualized via the chiPlot website [81].

### Identification of chromosomal inversions

Inversions were identified from JCVI-based gene synteny analysis using NGenomeSyn v1.42 [82] visualization, with evident funnel-shaped patterns considered inversion regions. Breakpoints were defined as 40 kb regions flanking the genes located at inversion boundaries in the HJD genome assembly. Enrichment analyses of repeats at inversion breakpoints were performed through permutation tests involving four categories of repetitive elements: LTR/Copia, LTR/Gypsy, DNA transposons, and segmental duplications. Specifically, centromeric and telomeric regions were masked genome-wide, and an in-house script was employed to conduct 1,000 permutation tests. Repeat lengths in equivalent genomic intervals provided expected values, with enrichment determined by the frequency and magnitude of observed values exceeding these expectations. In practice, we performed 1000 permutations and computed the p-value as p = (N − k) / N, where k is the number of permutations in which the observed repeat occupancy exceeded its expected value.

## Acknowledgements

This work was financially supported by State Key Laboratory for Tropical Crop Breeding and the start-up package from Agricultural Genomics Institute at Shenzhen, Chinese Academy of Agricultural Sciences.

## Author Contributions

Y.F.T. and Y.Y. conceived the project. C.Z.W., S.C.S., Y.Z.W. and S.P. performed bioinformatic analyses. Y.Z.W., Y.B.W., and T.D. conducted plant cultivation and experiments. C.Z.W. and Y.F.T. wrote the manuscript. E.M. D.J. supervised the work. All authors contributed to the article and approved the submitted version.

## Data availability

Raw sequencing reads and assembly results for HJD have been deposited under project PRJCA044301 at the National Genomics Data Center. Protein-coding g ene annotation files and additional assembly resources have been uploaded to t he online repository Figshare (URL: https://figshare.com/articles/dataset/cowpea_g enome_project/29878346). The scripts used for the analysis is available in a Git Hub repository at https://github.com/ChuanzhengWei/cowpea_T2T.

## Supplementary Data

Fig. S1 GenomeScope k-mer frequency profiles for HJD and FC6 sequencing data.

Fig. S2 Coverage landscape of HiFi and ONT reads mapped to the genome.

Fig. S3 BUSCO assessment results for HJD, FC6 and IT97K-499-35 genome assemblies using the Fabales reference database, with light blue indicating complete single-copy genes.

Fig. S4 K-mer frequency profiles generated by merqury for genome accuracy assessment.

Fig. S5 Physical and genetic distance comparison using IT97K-499-35 as the reference genome.

Fig. S6 Physical and genetic distance comparison using HJD as the reference genome, with black dashed lines indicating centromere positions.

Fig. S7 Reads mapping rate, genome coverage rate, and error rate obtained by aligning resequencing short reads to IT97K-499-35, HJD, and FC6 reference genomes. Letters a, b, c indicate statistical test results among groups (different letters denote significant differences at p < 0.05, identical letters indicate no significant difference).

Fig. S8 GO functional enrichment of centromeric genes.

Fig. S9 Genome synteny alignment between HJD and FC6.

Fig. S10 Mixed Ks plot centered on HJD generated by WGD. At Ks ∼0.73 there is a prominent Ks peak delineated by both the whole paranome and anchor pair Ks distributions, indicating the presence of an ancient WGD event. The corrected divergence times with other species (i.e., mung bean, common bean and FC6) were superimposed on the paralogous Ks distribution in vertical dashdot lines while in the legend it shows both the original (i.e. uncorrected) and corrected divergence times in the Ks timescale of the focal HJD.

Fig. S11 Dot plot of paralogous gene pairs from HJD self-alignment.

Fig. S12 GO enrichment results for contracted and expanded gene families, in the order of FC6 expanded genes, FC6 contracted genes, and HJD expanded genes.

Fig. S13 Distribution of major repeat monomers on chromosomes generated by TRASH software, with different colors indicating monomer lengths.

Fig. S14 Similarity of tandem arrays on soybean chromosomes, with clustered triangles representing TRAs and redder coloration indicating higher similarity.

Fig. S15 Similarity of tandem arrays on common bean chromosomes, with clustered triangles representing TRAs and redder coloration indicating higher similarity.

Fig. S16 Similarity of tandem arrays on mung bean chromosomes, with clustered triangles representing TRAs and redder coloration indicating higher similarity.

Fig. S17 Similarity of tandem arrays on cowpea chromosomes, with clustered triangles representing TRAs and redder coloration indicating higher similarity.

Fig. S18 Phylogenetic tree of selected CEN440 and CEN455 elements and corresponding sequence alignment of representative monomers.

Fig. S19 Phylogenetic tree and representative sequence alignment of Gm335, Vr215, Pv539 and HJD338.

Fig. S20 Synteny between cowpea and mung bean; central lines connect homologous chromosomes, colored blocks mark centromeres, gray lines link homologous genes, and green lines highlight genes within or adjacent to centromeres.

Fig. S21 Synteny between cowpea and common bean; central lines connect homologous chromosomes, colored blocks mark centromeres, gray lines link homologous genes, and green lines highlight genes within or adjacent to centromeres.

Fig. S22 Dot plots of inversions at shared breakpoints in cowpea versus Soybean, cowpea versus mung bean and cowpea versus common bean; blue dots represent orthologous gene pairs, gray shading indicates aligned regions.

Fig. S23 Copia, Gypsy and DNA transposons occupancy at inversion breakpoints estimated from 1000 random resampling iterations.

Table S1. Sequencing data generated from different platforms for assembly.

Table S2. Genome survey result statistics for HJD and FC6.

Table S3. Primary assembly statistics produced by hifiasm.

Table S4. Primary contig statistics produced by verkko and NextDenovo.

Table S5. Summary statistics of long-read mapping to the genome.

Table S6. Kmer accuracy statistics generated by merqury.

Table S7. Annotation statistics of annotated genes in databases.

Table S8. Distribution of telomeres in genome components.

Table S9. Distribution of complete CEN455 in genome components.

Table S10. Predicted locations of centromeres.

Table S11. Distribution of initial and improved inversion regions.

Table S12. Mapping rate of short reads aligned to the genome.

Table S13. Coverage rate of short reads aligned to the genome.

Table S14. Error rate of short reads aligned to the genome.

Table S15. GO functional enrichment of centromeric genes.

Table S16. Inversion positions and FST distribution in inversion regions.

Table S17. GO functional enrichment of genes within inversion regions.

Table S18. Species genome information used for phylogenetic tree construction.

Table S19. Paralogous gene pairs in the HJD genome.

Table S20. GO functional enrichment of expanded and contracted gene families.

Table S21. Positions of centromeric TRAs.

Table S22. Statistics of repeat sequence annotations.

Table S23. TE size and genome size of species within Phaseoleae.

Table S24. Composition of different repeat types within centromeres.

Table S25. Composition of different repeat element families within centromeres.

Table S26. Positions of SD regions in different genomes.

## References

1. Ehlers, J.D. Hall, A.E. Cowpea (*Vigna unguiculata* L. Walp.). Field Crops Res. 1997;53:187–204.

2. Chen, H., Chen, H., Hu, L. et al. Genetic diversity and a population structure analysis of accessions in the Chinese cowpea [*Vigna unguiculata* (L.) Walp.] germplasm collection. Crop J. 2017;5:363–372.

3. Chipeta, M.M., Kafwambira, J. Yohane, E. Cowpea genetic diversity, population structure and genome-wide association studies in Malawi: insights for breeding programs. Front Plant Sci. 2025;15:1461631.

4. Boukar, O., Fatokun, C.A., Huynh, B.L. et al. Genomic Tools in Cowpea Breeding Programs: Status and Perspectives. Front Plant Sci. 2016;7:757.

5. Pasquet, R.S. Allozyme diversity of cultivated cowpea *Vigna unguiculata* (L.) Walp. Theor Appl Genet. 2000;101:211–219.

6. Kouakou, A.G., Ogundapo, A., Smale, M., Jamora, N., Manda, J. Abberton, M. IITA’s genebank, cowpea diversity on farms, and farmers’ welfare in Nigeria. CABI Agri Biosci. 2022;3:14.

7. Fang, J., Chao, C.-C.T., Roberts, P.A. Ehlers, J.D. Genetic diversity of cowpea [*Vigna unguiculata* (L.) Walp.] in four West African and USA breeding programs as determined by AFLP analysis. Genet Resour Crop Evol. 2007;54:1197–1209.

8. Xu, P., Wu, X., Wang, B. et al. Development and polymorphism of *Vigna unguiculata* ssp. *unguiculata* microsatellite markers used for phylogenetic analysis in asparagus bean (*Vigna unguiculata* ssp. *sesquipedialis* (L.) Verdc.). Mol Breed. 2010;25:675–684.

9. Jayathilake, C., Visvanathan, R., Deen, A. et al. Cowpea: an overview on its nutritional facts and health benefits. J Sci Food Agric. 2018;98:4793–4806.

10. Affrifah, N.S., Phillips, R.D. Saalia, F.K. Cowpeas: Nutritional profile, processing methods and products—A review. Legume Sci. 2022;4:e131.

11. Della Coletta, R., Qiu, Y., Ou, S., Hufford, M.B. Hirsch, C.N. How the pan-genome is changing crop genomics and improvement. Genome Bio. 2021;22:3.

12. Muñoz-Amatriaín, M., Mirebrahim, H., Xu, P. et al. Genome resources for climate-resilient cowpea, an essential crop for food security. Plant J. 2017;89:1042–1054.

13. Lonardi, S., Muñoz-Amatriaín, M., Liang, Q. et al. The genome of cowpea (*Vigna unguiculata* [L.] Walp.). Plant J. 2019;98:767–782.

14. Xia, Q., Pan, L., Zhang, R. et al. The genome assembly of asparagus bean, Vigna unguiculata ssp. sesquipedialis. Sci Data. 2019;6:124.

15. Yang, Y., Wu, Z., Wu, Z. et al. A near-complete assembly of asparagus bean provides insights into anthocyanin accumulation in pods. Plant Biotechnol J. 2023;21:2473–2489.

16. Kajita, T., Ohashi, H., Tateishi, Y., Bailey, C.D. Doyle, J.J. rbcL and Legume Phylogeny, with Particular Reference to Phaseoleae, Millettieae, and Allies. Syst Bot. 2001;26:515–536.

17. Lackey, J.A. Chromosome numbers in the Phaseoleae (Fabaceae: Faboideae) and their relation to taxonomy. Am J Bot. 1980;67:595–602.

18. Vasconcelos, E.V., de Andrade Fonsêca, A.F., Pedrosa-Harand, A. et al. Intra- and interchromosomal rearrangements between cowpea [*Vigna unguiculata* (L.) Walp.] and common bean (*Phaseolus vulgaris* L.) revealed by BAC-FISH. Chromosome Res. 2015;23:253–266.

19. de Oliveira Bustamante, F., do Nascimento, T.H., Montenegro, C., et al. Oligo-FISH barcode in beans: a new chromosome identification system. Theor Appl Genet. 2021;134:3675–3686.

20. Oliveira, A.R.d.S., Martins, L.d.V., Bustamante, F.d.O. et al. Breaks of macrosynteny and collinearity among moth bean (*Vigna aconitifolia*), cowpea (*V. unguiculata*), and common bean (*Phaseolus vulgaris*). Chromosome Res. 2020;28:293–306.

21. Cleveland, D.W., Mao, Y. Sullivan, K.F. Centromeres and Kinetochores: From Epigenetics to Mitotic Checkpoint Signaling. Cell. 2003;112:407–421.

22. do Vale Martins, L., de Oliveira Bustamante, F., da Silva Oliveira, A.R. et al. BAC- and oligo-FISH mapping reveals chromosome evolution among *Vigna angularis*, V. unguiculata, and Phaseolus vulgaris. Chromosoma. 2021;130:133–147.

23. Montenegro, C., do Vale Martins, L., Bustamante, F.d.O., Brasileiro-Vidal, A.C. Pedrosa-Harand, A. Comparative cytogenomics reveals genome reshuffling and centromere repositioning in the legume tribe Phaseoleae. Chromosome Res. 2022;30:477–492.

24. Wang, L., Zhang, M., Li, M., Jiang, X., Jiao, W. Song, Q. A telomere-to-telomere gap-free assembly of soybean genome. Mol Plant. 2023;16:1711–1714.

25. Wang, Y., Hao, X., Chen, C. et al. Telomere-to-telomere genome of common bean (Phaseolus vulgaris L., YP4). GigaScience. 2025;14:giaf001.

26. Zhao, B., Zhang, H., Zhao, Q. et al. Gap-free genome assembly and metabolomics analysis of common bean provide insights into genomic characteristics and metabolic determinants of seed coat pigmentation. J Genet Genomics. 2025;52:852–855.

27. Jia, K.-H., Li, G., Wang, L. et al. Telomere-to-telomere, gap-free genome of mung bean (*Vigna radiata*) provides insights into domestication under structural variation. Hortic Res. 2025;12:uhae337.

28. Liu, J., Li, Q., Hu, Y. et al. The complete telomere-to-telomere sequence of a mouse genome. Science. 2024;386:1141–1146.

29. Naish, M. Henderson, I.R. The structure, function, and evolution of plant centromeres. Genome Res. 2024;34:161–178.

30. Pan, L., Liu, M., Kang, Y. et al. Comprehensive genomic analyses of *Vigna unguiculata* provide insights into population differentiation and the genetic basis of key agricultural traits. Plant Biotechnol J. 2023;21:1426–1439.

31. Wellenreuther, M. Bernatchez, L. Eco-Evolutionary Genomics of Chromosomal Inversions. Trends Ecol Evol. 2018;33:427–440.

32. Wang, J., Sun, P., Li, Y. et al. Hierarchically Aligning 10 Legume Genomes Establishes a Family-Level Genomics Platform. Plant Physiol. 2017;174:284–300.

33. Schmutz, J., Cannon, S.B., Schlueter, J. et al. Genome sequence of the palaeopolyploid soybean. Nature. 2010;463:178–183.

34. Young, N.D., Debellé, F., Oldroyd, G.E.D. et al. The Medicago genome provides insight into the evolution of rhizobial symbioses. Nature. 2011;480:520–524.

35. Shang, L., He, W., Wang, T. et al. A complete assembly of the rice Nipponbare reference genome. Mol Plant. 2023;16:1232–1236.

36. Chen, J., Wang, Z., Tan, K. et al. A complete telomere-to-telomere assembly of the maize genome. Nat Genet. 2023;55:1221–1231.

37. Wei, C., Gao, L., Xiao, R. et al. Complete telomere-to-telomere assemblies of two sorghum genomes to guide biological discovery. iMeta. 2024;3:e193.

38. Kumon, T. Lampson, M.A. Evolution of eukaryotic centromeres by drive and suppression of selfish genetic elements. Semin Cell Dev Biol. 2022;128:51–60.

39. Hu, G., Wang, Z., Tian, Z. et al. A telomere-to-telomere genome assembly of cotton provides insights into centromere evolution and short-season adaptation. Nat Genet. 2025;57:1031–1043.

40. Jin, X., Du, H., Chen, M., Zheng, X., He, Y. Zhu, A. A fully phased octoploid strawberry genome reveals the evolutionary dynamism of centromeric satellites. Genome Biol. 2025;26:17.

41. Courret, C., Hemmer, L.W., Wei, X. et al. Turnover of retroelements and satellite DNA drives centromere reorganization over short evolutionary timescales in Drosophila. PLoS Biol. 2024;22:e3002911.

42. Zhao, J., Xie, Y., Kong, C. et al. Centromere repositioning and shifts in wheat evolution. Plant Commun. 2023;4:100556.

43. Wang, Y., Zhao, L., Wang, D. et al. Four near-complete genome assemblies reveal the landscape and evolution of centromeres in Salicaceae. Genome Biol. 2025;26:111.

44. Huang, G., Wang, X., Liu, C. et al. Genomic Variation Underpins Genetic Divergence and Differing Salt Resilience in *Sesbania bispinosa*. Adv Sci. 2025;e02600.

45. Blumer, L.M., Burskaia, V., Artiushin, I. et al. Introgression dynamics of sex-linked chromosomal inversions shape the Malawi cichlid radiation. Science. 2025;388:eadr9961.

46. Porebski, S., Bailey, L.G. Baum, B.R. Modification of a CTAB DNA extraction protocol for plants containing high polysaccharide and polyphenol components. Plant Mol Biol Rep. 1997;15:8–15.

47. Rao, Suhas S.P., Huntley, Miriam H., Durand, Neva C. et al. A 3D Map of the Human Genome at Kilobase Resolution Reveals Principles of Chromatin Looping. Cell. 2014;159:1665–1680.

48. Marçais, G. Kingsford, C. A fast, lock-free approach for efficient parallel counting of occurrences of k-mers. Bioinformatics. 2011;27:764–770.

49. Ranallo-Benavidez, T.R., Jaron, K.S. Schatz, M.C. GenomeScope 2.0 and Smudgeplot for reference-free profiling of polyploid genomes. Nat Commun. 2020;11:1432.

50. Cheng, H., Jarvis, E.D., Fedrigo, O. et al. Haplotype-resolved assembly of diploid genomes without parental data. Nat Biotechnol. 2022;40:1332–1335.

51. Li, H. Aligning sequence reads, clone sequences and assembly contigs with BWA-MEM. arXiv preprint. 2013;1303.3997.

52. Zeng, X., Yi, Z., Zhang, X. et al. Chromosome-level scaffolding of haplotype-resolved assemblies using Hi-C data without reference genomes. Nat Plants. 2024;10:1184–1200.

53. Rautiainen, M., Nurk, S., Walenz, B.P. et al. Telomere-to-telomere assembly of diploid chromosomes with Verkko. Nat Biotechnol. 2023;41:1474–1482.

54. Hu, J., Wang, Z., Sun, Z. et al. NextDenovo: an efficient error correction and accurate assembly tool for noisy long reads. Genome Biol. 2024;25:107.

55. Jain, C., Rhie, A., Hansen, N.F., Koren, S. Phillippy, A.M. Long-read mapping to repetitive reference sequences using Winnowmap2. Nature Methods. 2022;19:705–710.

56. Manni, M., Berkeley, M.R., Seppey, M., Simão, F.A. Zdobnov, E.M. BUSCO Update: Novel and Streamlined Workflows along with Broader and Deeper Phylogenetic Coverage for Scoring of Eukaryotic, Prokaryotic, and Viral Genomes. Mol Biol Evol. 2021;38:4647–4654.

57. Li, H. Minimap2: pairwise alignment for nucleotide sequences. Bioinformatics. 2018;34:3094–3100.

58. Yu, H., Shi, C., He, W., Li, F. Ouyang, B. PanDepth, an ultrafast and efficient genomic tool for coverage calculation. Briefings Bioinf. 2024;25:bbae197.

59. Rhie, A., Walenz, B.P., Koren, S. Phillippy, A.M. Merqury: reference-free quality, completeness, and phasing assessment for genome assemblies. Genome Biol. 2020;21:245.

60. Hu, K., Ni, P., Xu, M. et al. HiTE: a fast and accurate dynamic boundary adjustment approach for full-length transposable element detection and annotation. Nat Commun. 2024;15:5573.

61. Zhang, R.-G., Li, G.-Y., Wang, X.-L. et al. TEsorter: An accurate and fast method to classify LTR-retrotransposons in plant genomes. Hortic Res. 2022;9:uhac017.

62. Išerić, H., Alkan, C., Hach, F. Numanagić, I. Fast characterization of segmental duplication structure in multiple genome assemblies. Algorithms Mol Biol. 2022;17:4.

63. Kim, D., Paggi, J.M., Park, C., Bennett, C. Salzberg, S.L. Graph-based genome alignment and genotyping with HISAT2 and HISAT-genotype. Nat Biotechnol. 2019;37:907–915.

64. Gabriel, L., Brůna, T., Hoff, K.J. et al. BRAKER3: Fully automated genome annotation using RNA-seq and protein evidence with GeneMark-ETP, AUGUSTUS, and TSEBRA. Genome Res. 2024;34:769–777.

65. Pertea, M., Pertea, G.M., Antonescu, C.M., Chang, T.-C., Mendell, J.T. Salzberg, S.L. StringTie enables improved reconstruction of a transcriptome from RNA-seq reads. Nat Biotechnol. 2015;33:290–295.

66. Haas, B.J., Delcher, A.L., Mount, S.M. et al. Improving the Arabidopsis genome annotation using maximal transcript alignment assemblies. Nucleic Acids Res. 2003;31:5654–5666.

67. Buchfink, B., Xie, C. Huson, D.H. Fast and sensitive protein alignment using DIAMOND. Nat Methods. 2015;12:59–60.

68. Vollger, M.R., Kerpedjiev, P., Phillippy, A.M. Eichler, E.E. StainedGlass: interactive visualization of massive tandem repeat structures with identity heatmaps. Bioinformatics. 2022;38:2049–2051.

69. Wlodzimierz, P., Hong, M. Henderson, I.R. TRASH: Tandem Repeat Annotation and Structural Hierarchy. Bioinformatics. 2023;39:btad308.

70. Van der Auwera, G.A., Carneiro, M.O., Hartl, C. et al. From FastQ Data to High-Confidence Variant Calls: The Genome Analysis Toolkit Best Practices Pipeline. Curr Protoc Bioinf. 2013;43:11.10.1-11.10.33.

71. Chang, C.C., Chow, C.C., Tellier, L.C.A.M., Vattikuti, S., Purcell, S.M. Lee, J.J. Second-generation PLINK: rising to the challenge of larger and richer datasets. GigaScience. 2015;4:s13742–015-0047-8.

72. Robinson, J.T., Thorvaldsdóttir, H., Winckler, W. et al. Integrative genomics viewer. Nat Biotechnol. 2011;29:24–26.

73. Goel, M., Sun, H., Jiao, W.-B. Schneeberger, K. SyRI: finding genomic rearrangements and local sequence differences from whole-genome assemblies. Genome Biol. 2019;20:277.

74. Goel, M. Schneeberger, K. plotsr: visualizing structural similarities and rearrangements between multiple genomes. Bioinformatics. 2022;38:2922–2926.

75. Emms, D.M. Kelly, S. OrthoFinder: phylogenetic orthology inference for comparative genomics. Genome Biol. 2019;20:238.

76. Yang, Z. PAML 4: Phylogenetic Analysis by Maximum Likelihood. Mol Biol Evol. 2007;24:1586–1591.

77. Mendes, F.K., Vanderpool, D., Fulton, B. Hahn, M.W. CAFE 5 models variation in evolutionary rates among gene families. Bioinformatics. 2021;36:5516–5518.

78. Wu, T., Hu, E., Xu, S. et al. clusterProfiler 4.0: A universal enrichment tool for interpreting omics data. Innovation. 2021;2:100141.

79. Tang, H., Krishnakumar, V., Zeng, X. et al. JCVI: A versatile toolkit for comparative genomics analysis. iMeta. 2024;3:e211.

80. Chen, H., Zwaenepoel, A. Van de Peer, Y. wgd v2: a suite of tools to uncover and date ancient polyploidy and whole-genome duplication. Bioinformatics. 2024;40:btae272.

81. Xie, J., Chen, Y., Cai, G., Cai, R., Hu, Z. Wang, H. Tree Visualization By One Table (tvBOT): a web application for visualizing, modifying and annotating phylogenetic trees. Nucleic Acids Res. 2023;51:W587–W592.

82. He, W., Yang, J., Jing, Y., Xu, L., Yu, K. Fang, X. NGenomeSyn: an easy-to-use and flexible tool for publication-ready visualization of syntenic relationships across multiple genomes. Bioinformatics. 2023;39:btad121.

