## supplemental figure for "Complete telomere-to-telomere genomes of cowpea reveal insights into centromere evolution in Phaseoleae"

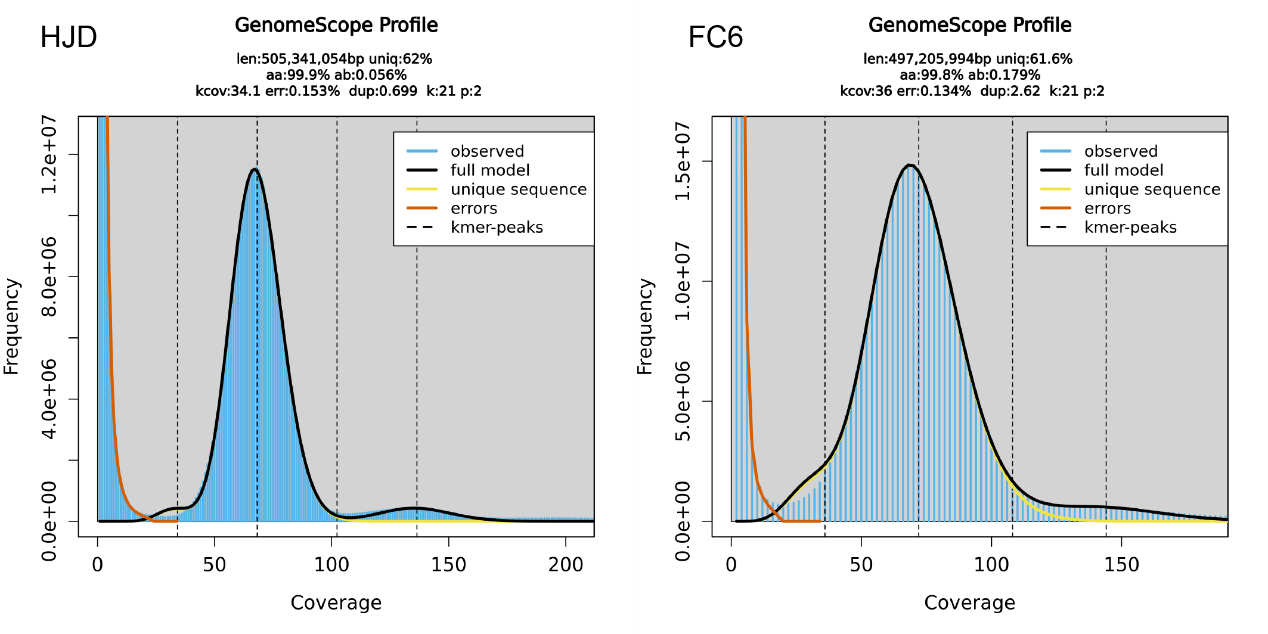

Fig. S1 GenomeScope k-mer frequency profiles for HJD and FC6 sequencing data.

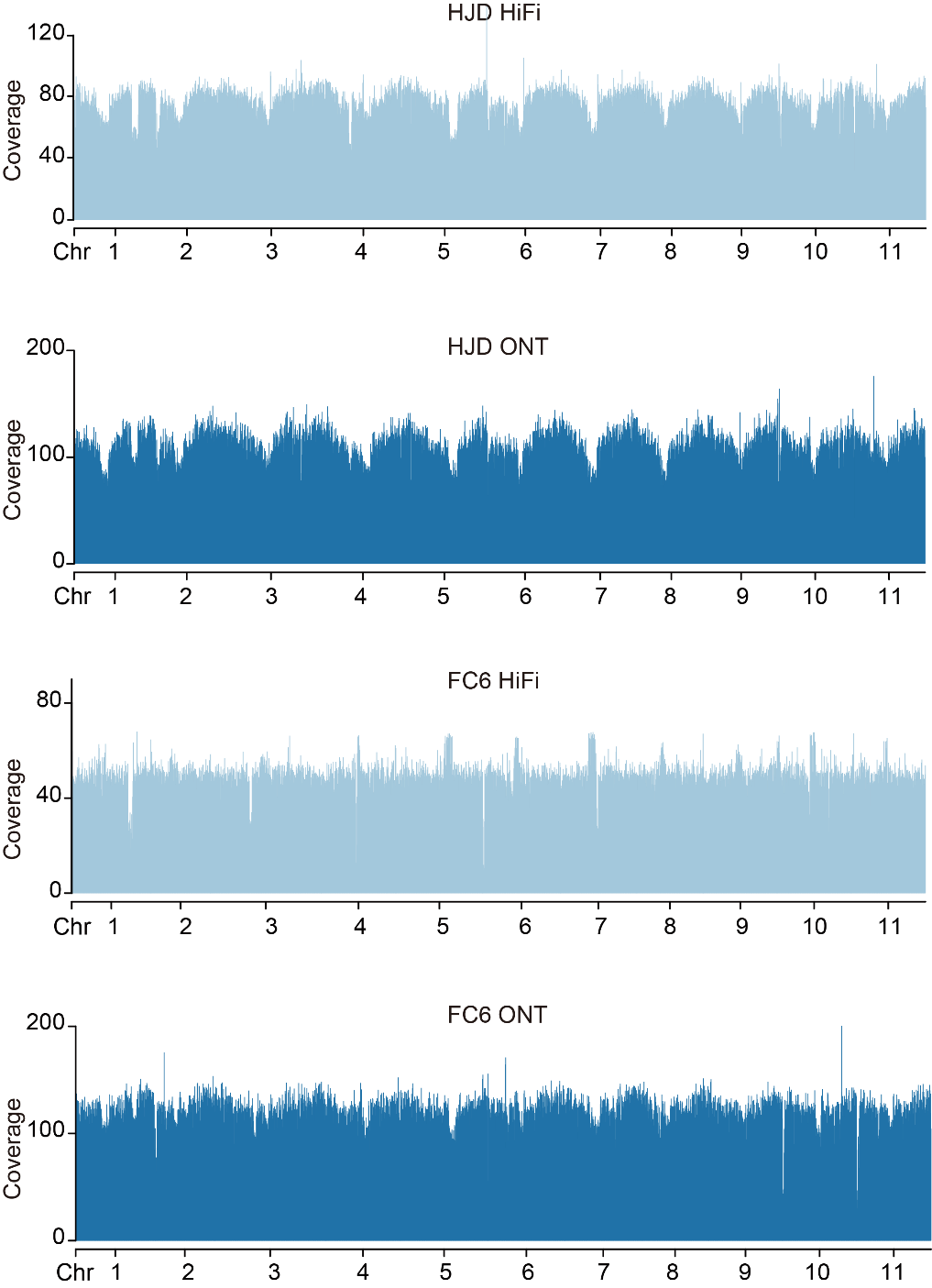

Fig. S2 Coverage landscape of HiFi and ONT reads mapped to the genome.

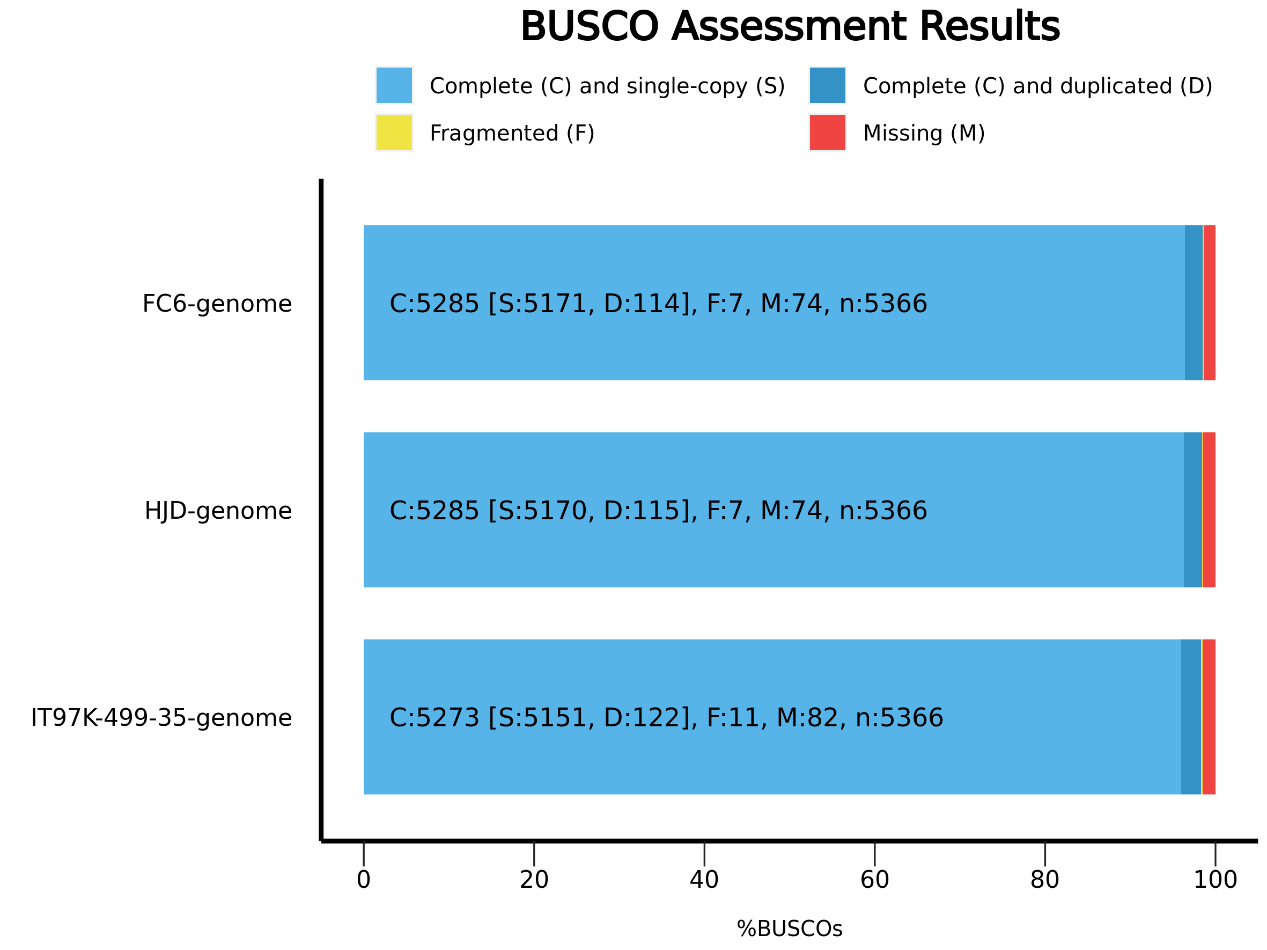

Fig. S3 BUSCO assessment results for HJD, FC6 and IT97K-499-35 genome assemblies using the Fabales reference database, with light blue indicating complete single-copy genes.

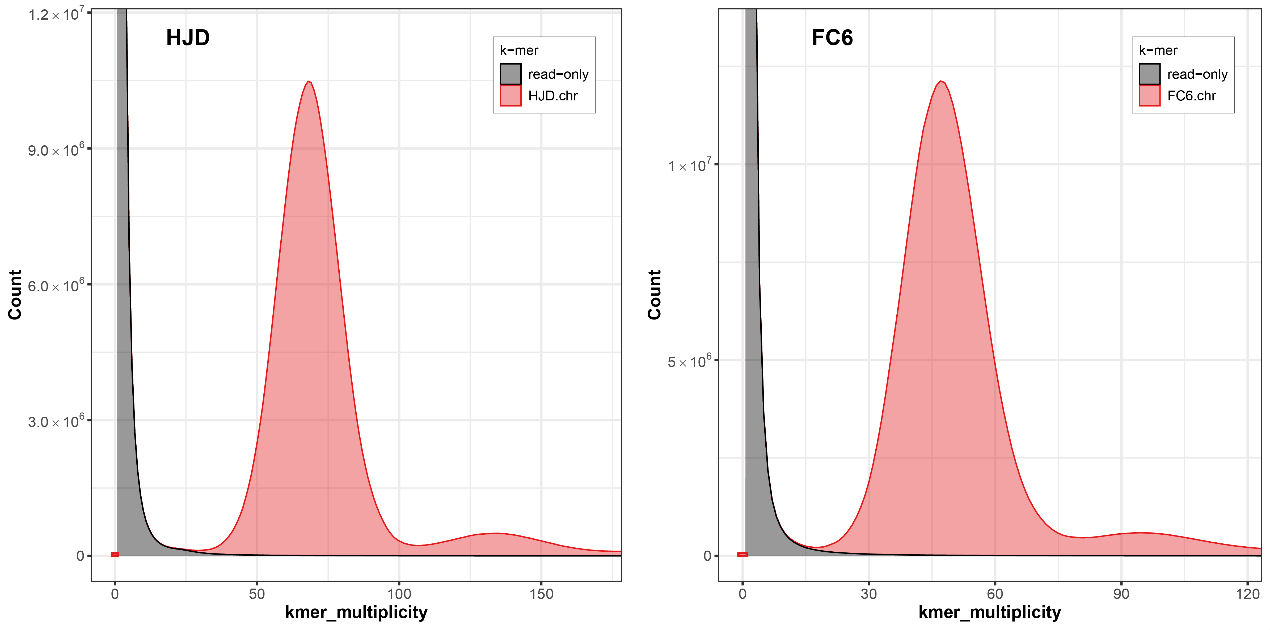

Fig. S4 K-mer frequency profiles generated by merqury for genome accuracy assessment.

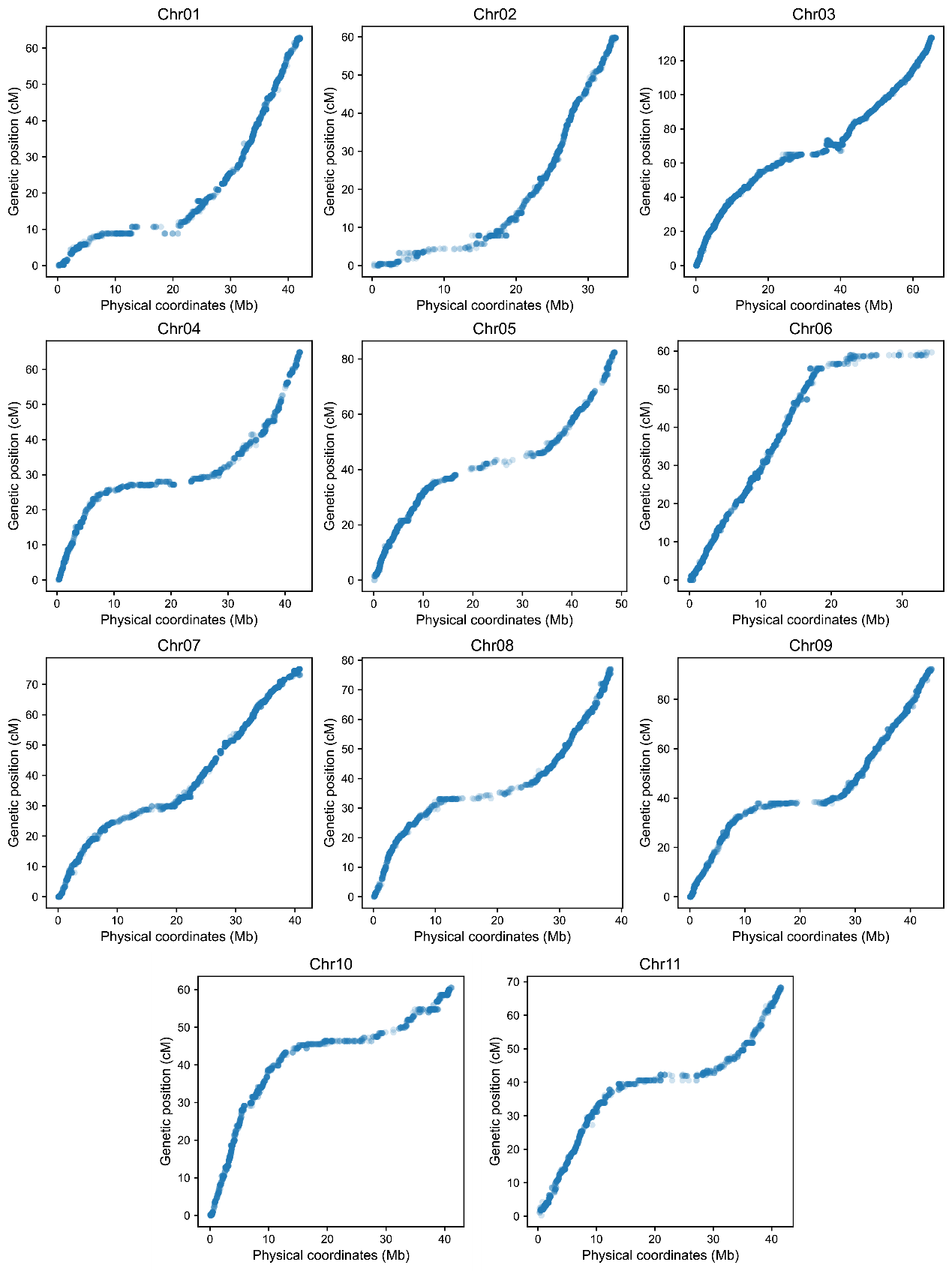

Fig. S5 Physical and genetic distance comparison using IT97K-499-35 as the reference genome.

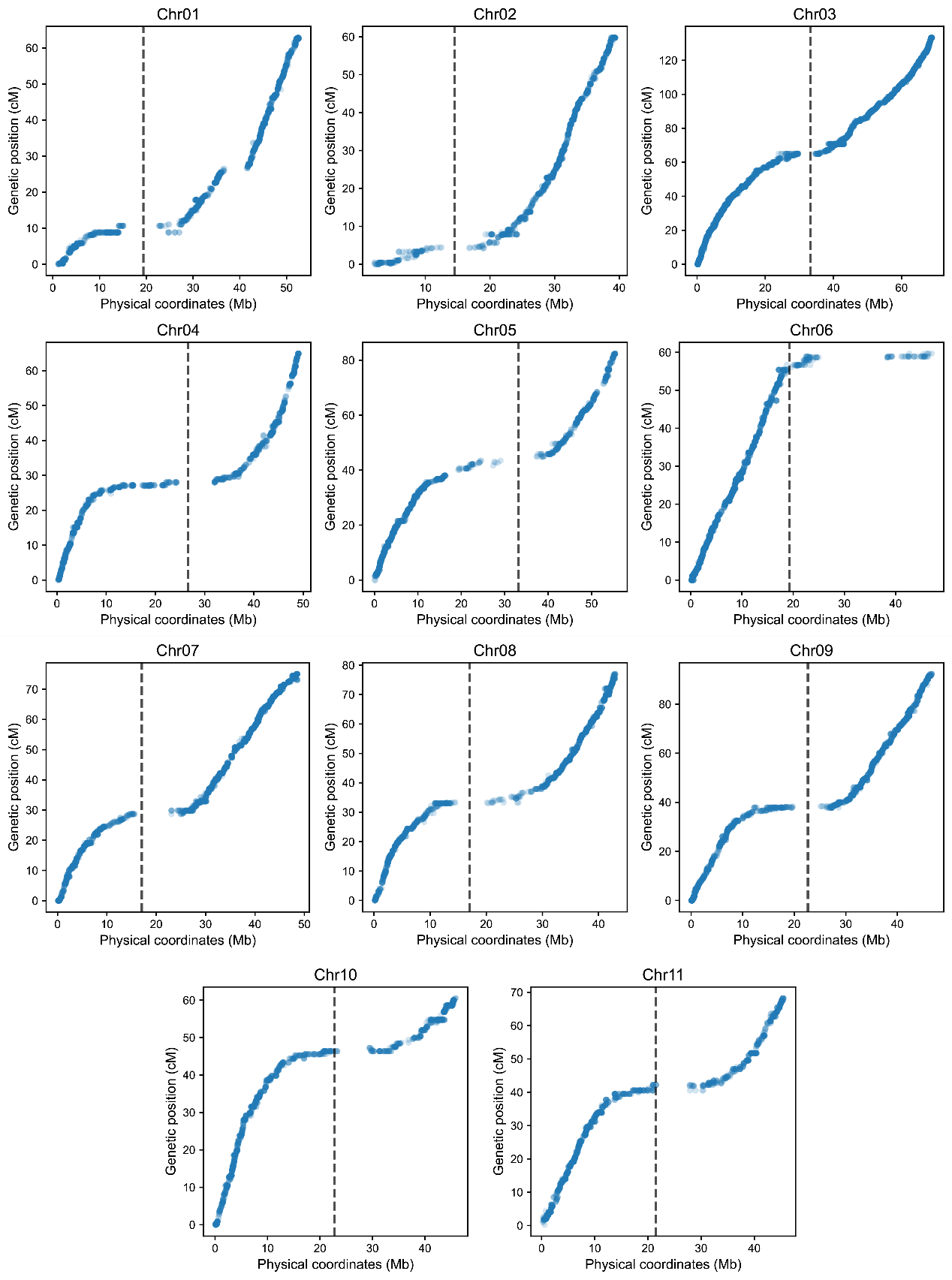

Fig. S6 Physical and genetic distance comparison using HJD as the reference genome, with black dashed lines indicating centromere positions.

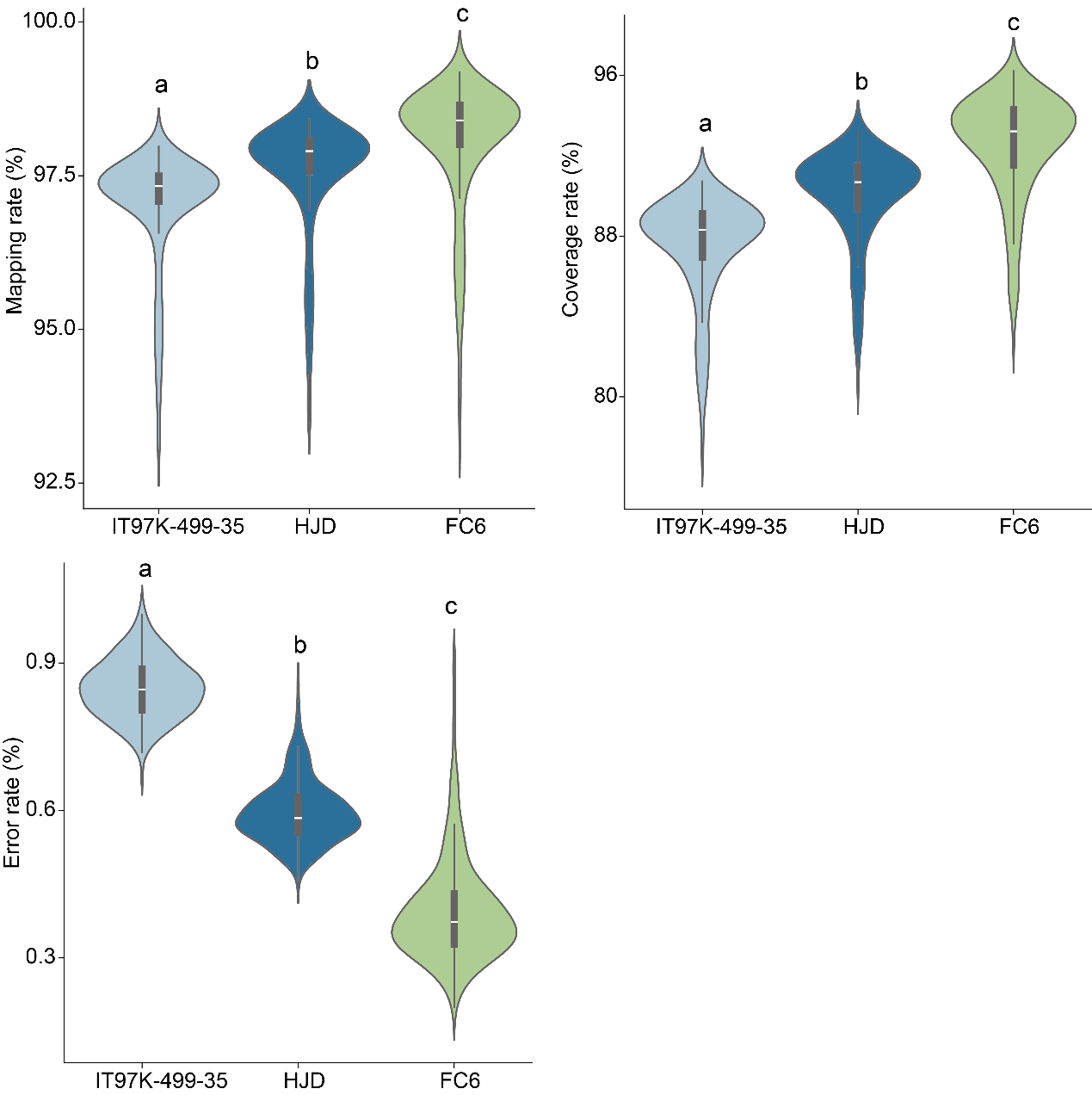

Fig. S7 Reads mapping rate, genome coverage rate, and error rate obtained by aligning resequencing short reads to IT97K-499-35, HJD, and FC6 reference genomes. Letters a, b, c indicate statistical test results among groups (different letters denote significant differences at p < 0.05, identical letters indicate no significant difference).

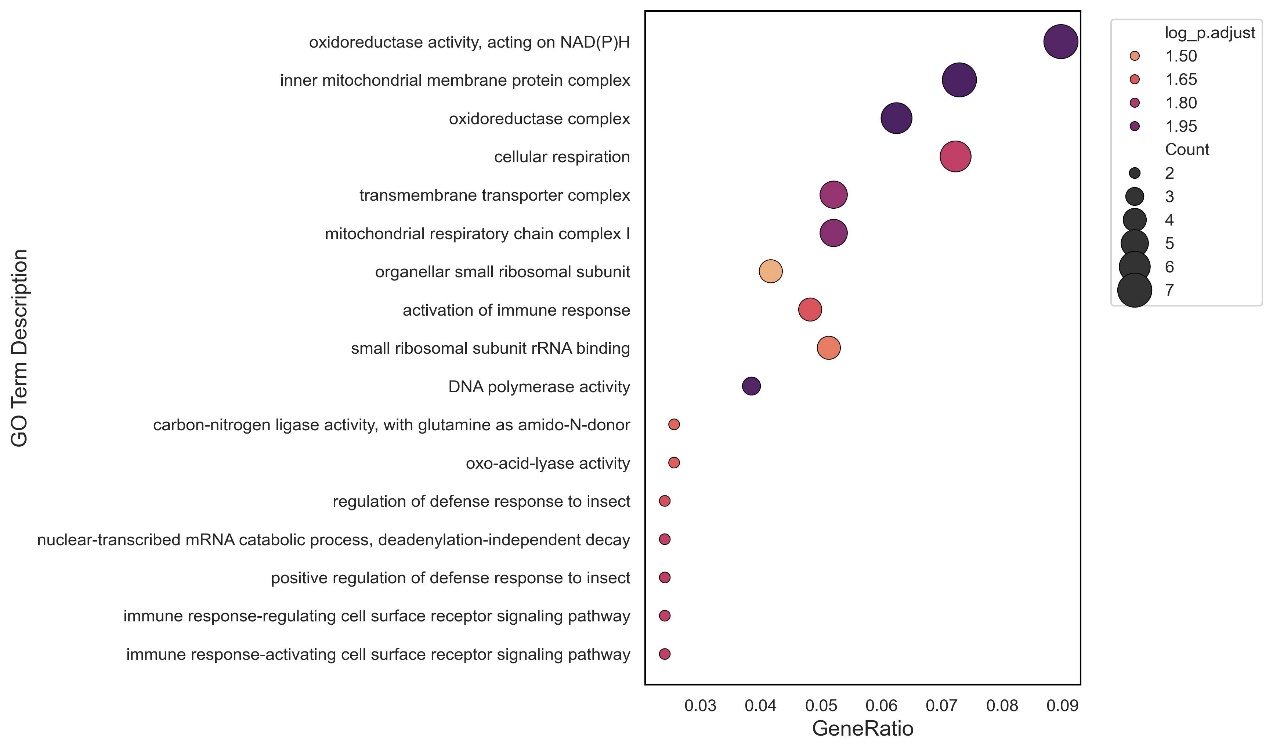

Fig. S8 GO functional enrichment of centromeric genes.

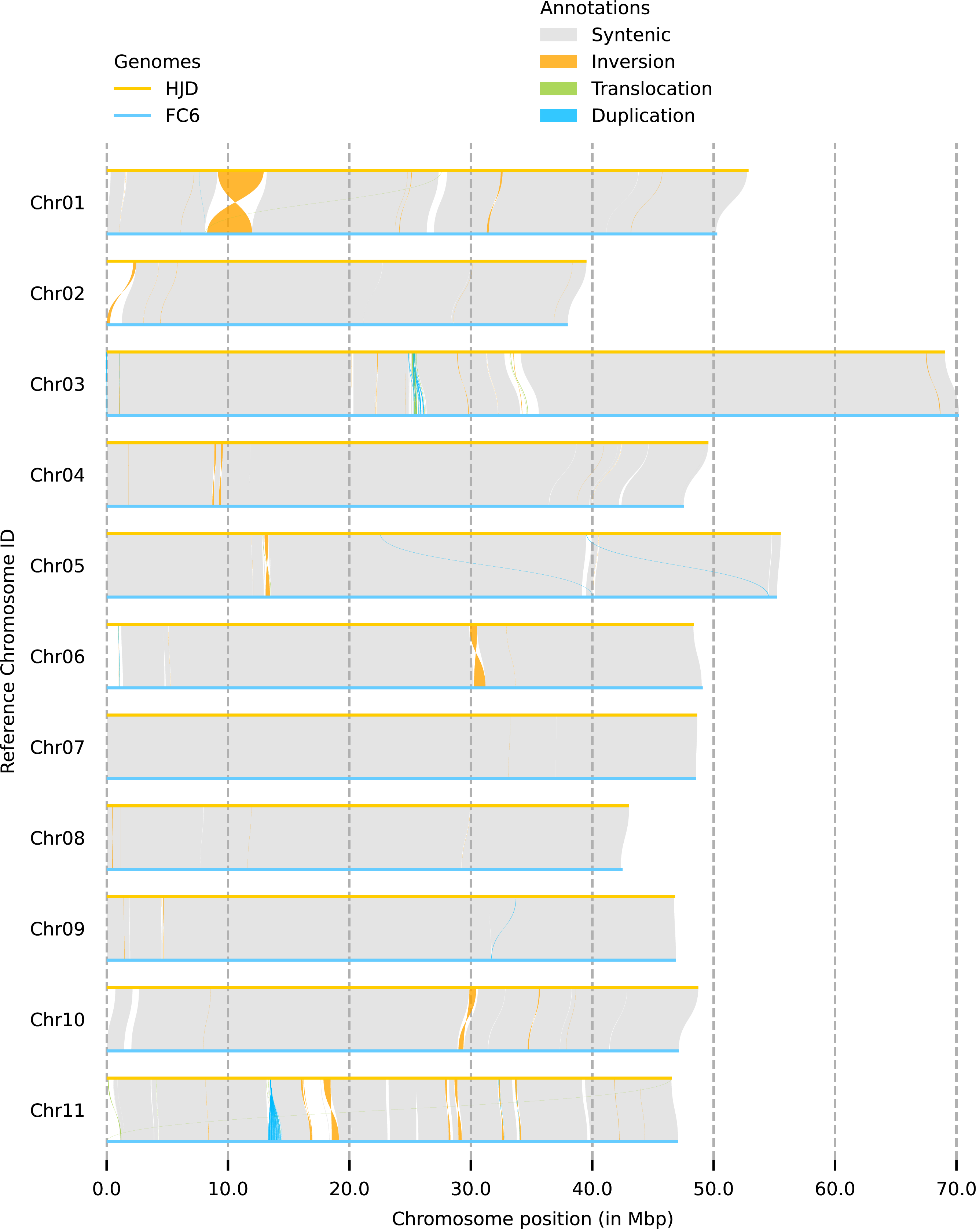

Fig. S9 Genome synteny alignment between HJD and FC6.

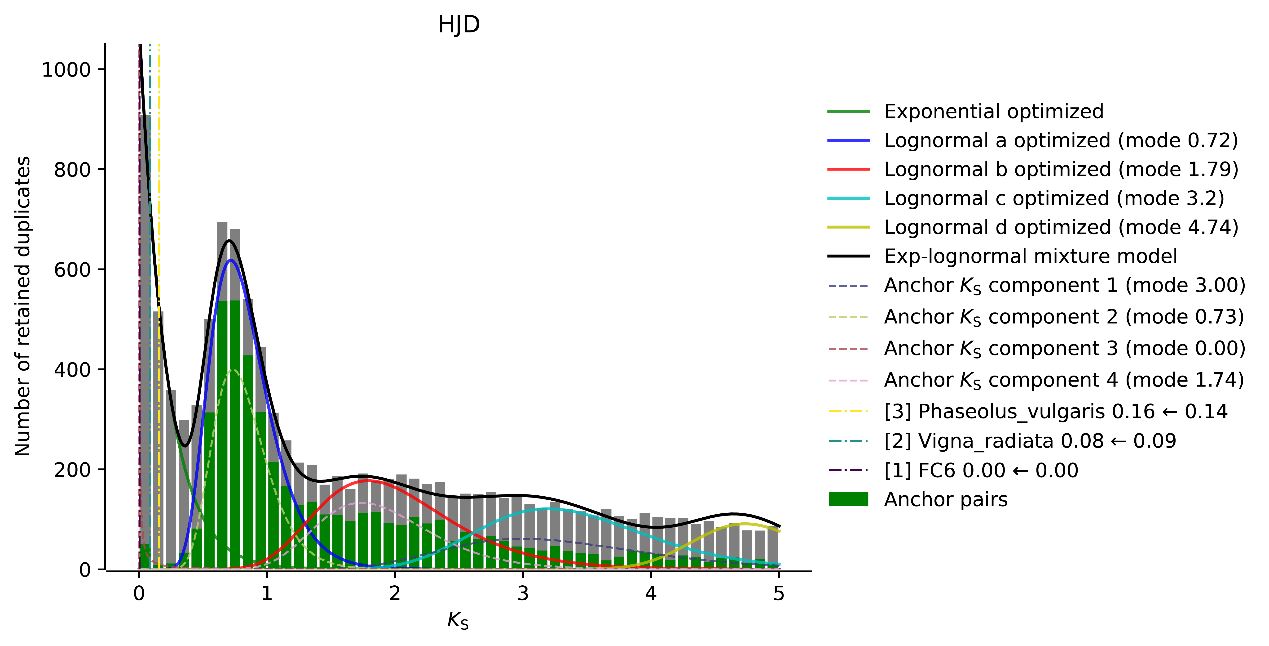

Fig. S10 Mixed Ks plot centered on HJD generated by WGD. At Ks ~0.73 there is a prominent Ks peak delineated by both the whole paranome and anchor pair Ks distributions, indicating the presence of an ancient WGD event. The corrected divergence times with other species (i.e., mung bean, common bean and FC6) were superimposed on the paralogous Ks distribution in vertical dashdot lines while in the legend it shows both the original (i.e. uncorrected) and corrected divergence times in the Ks timescale of the focal HJD.

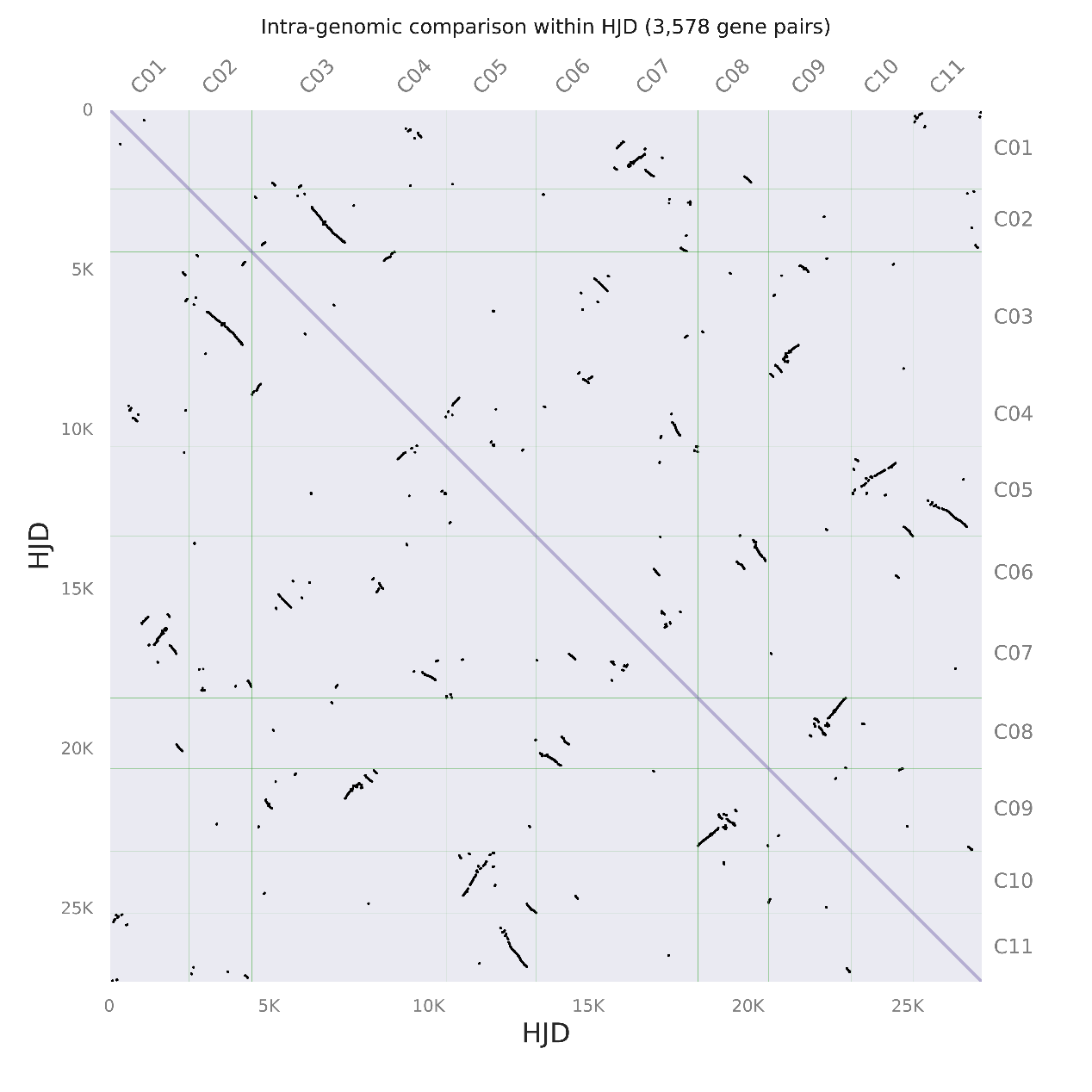

Fig. S11 Dot plot of paralogous gene pairs from HJD self-alignment.

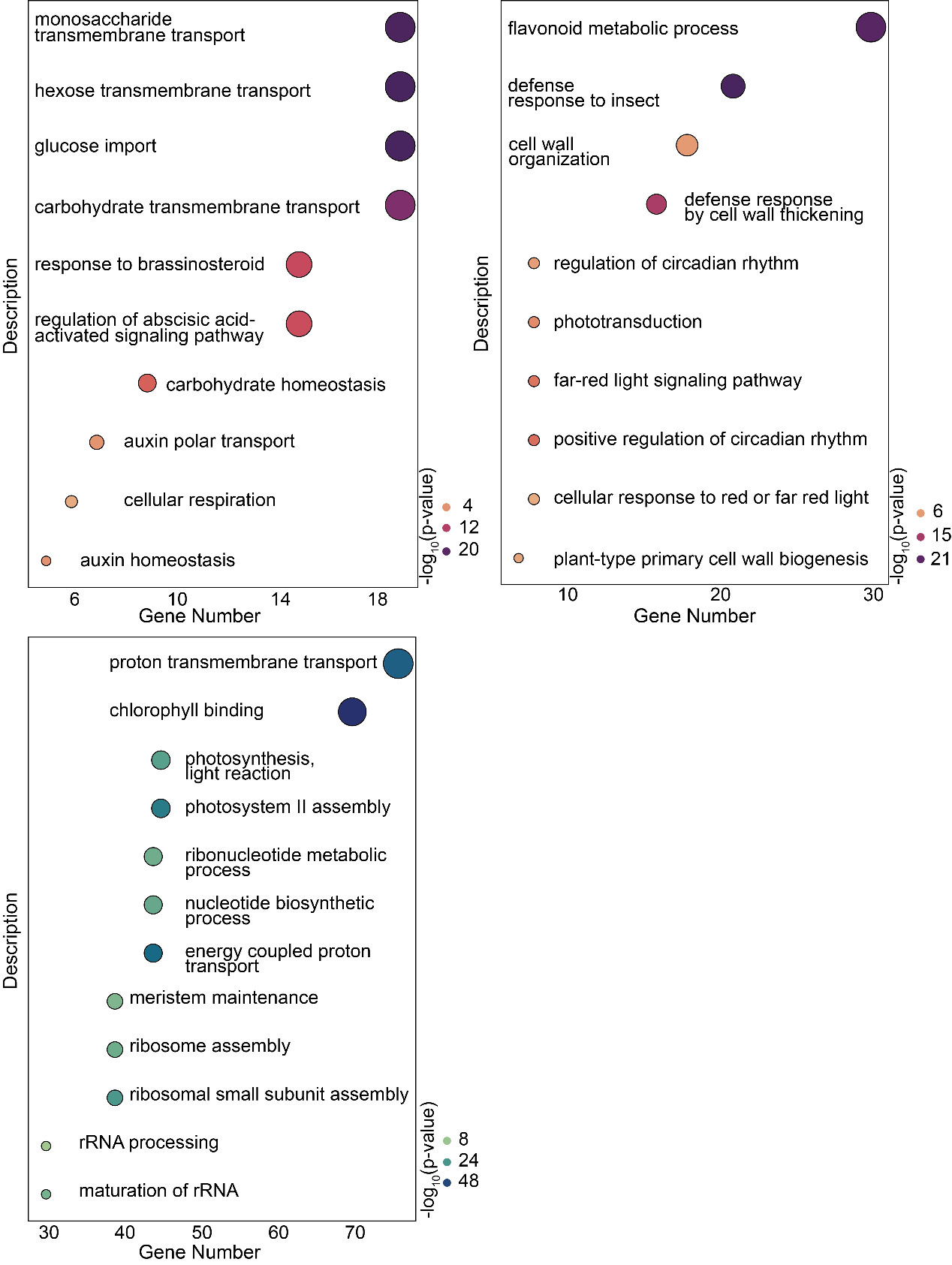

Fig. S12 GO enrichment results for contracted and expanded gene families, in the order of FC6 expanded genes, FC6 contracted genes, and HJD expanded genes.

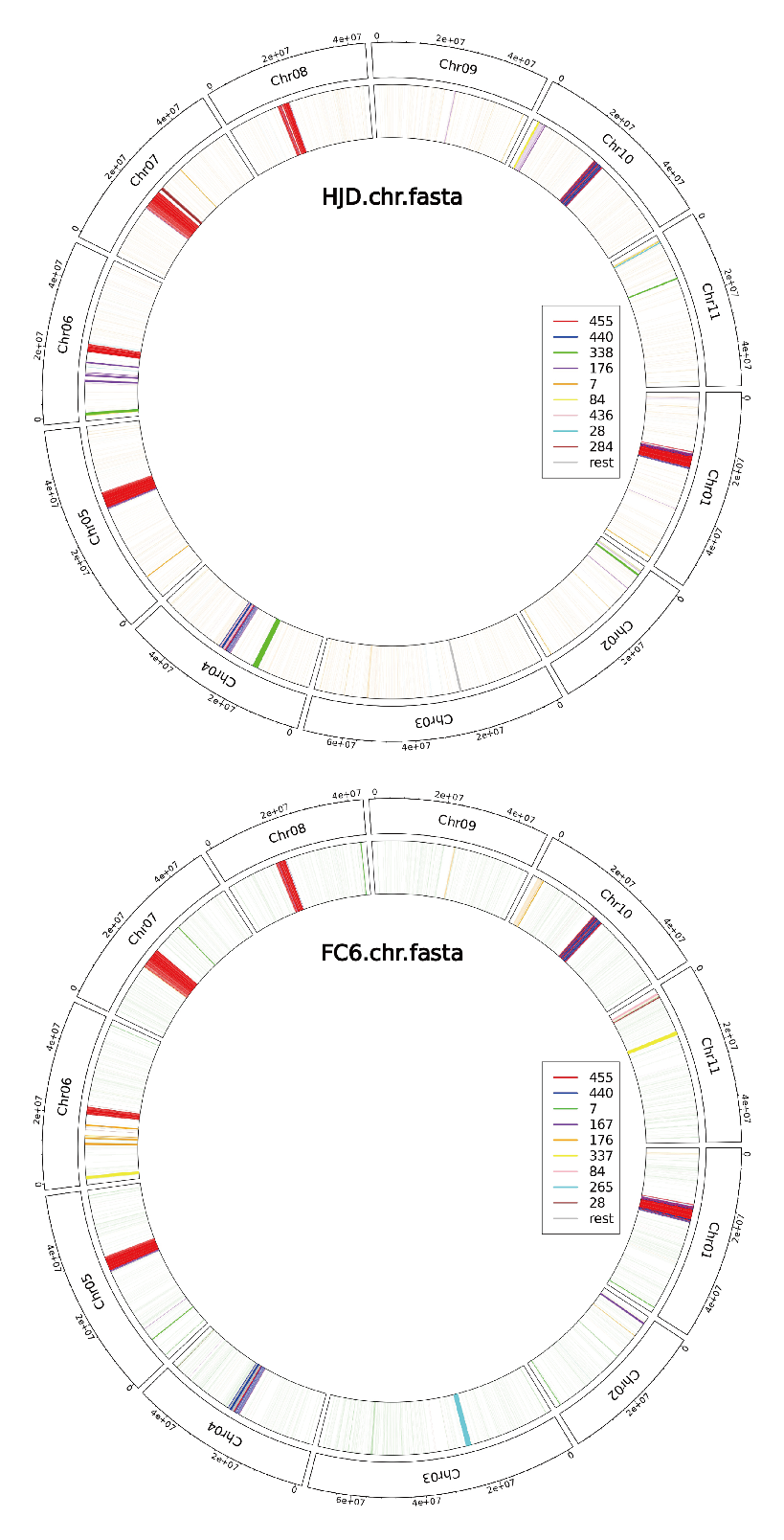

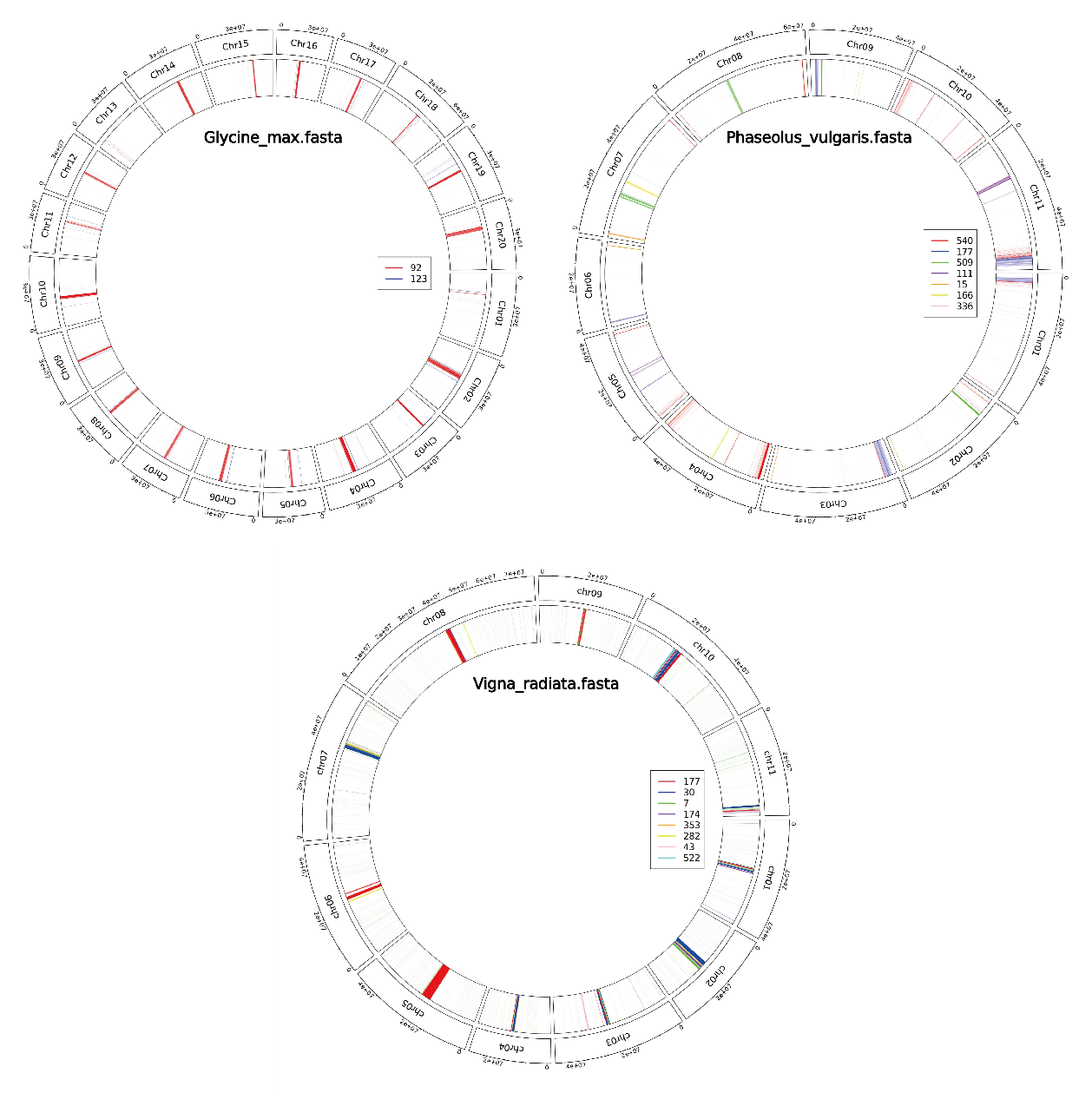

Fig. S13 Distribution of major repeat monomers on chromosomes generated by TRASH software, with different colors indicating monomer lengths.

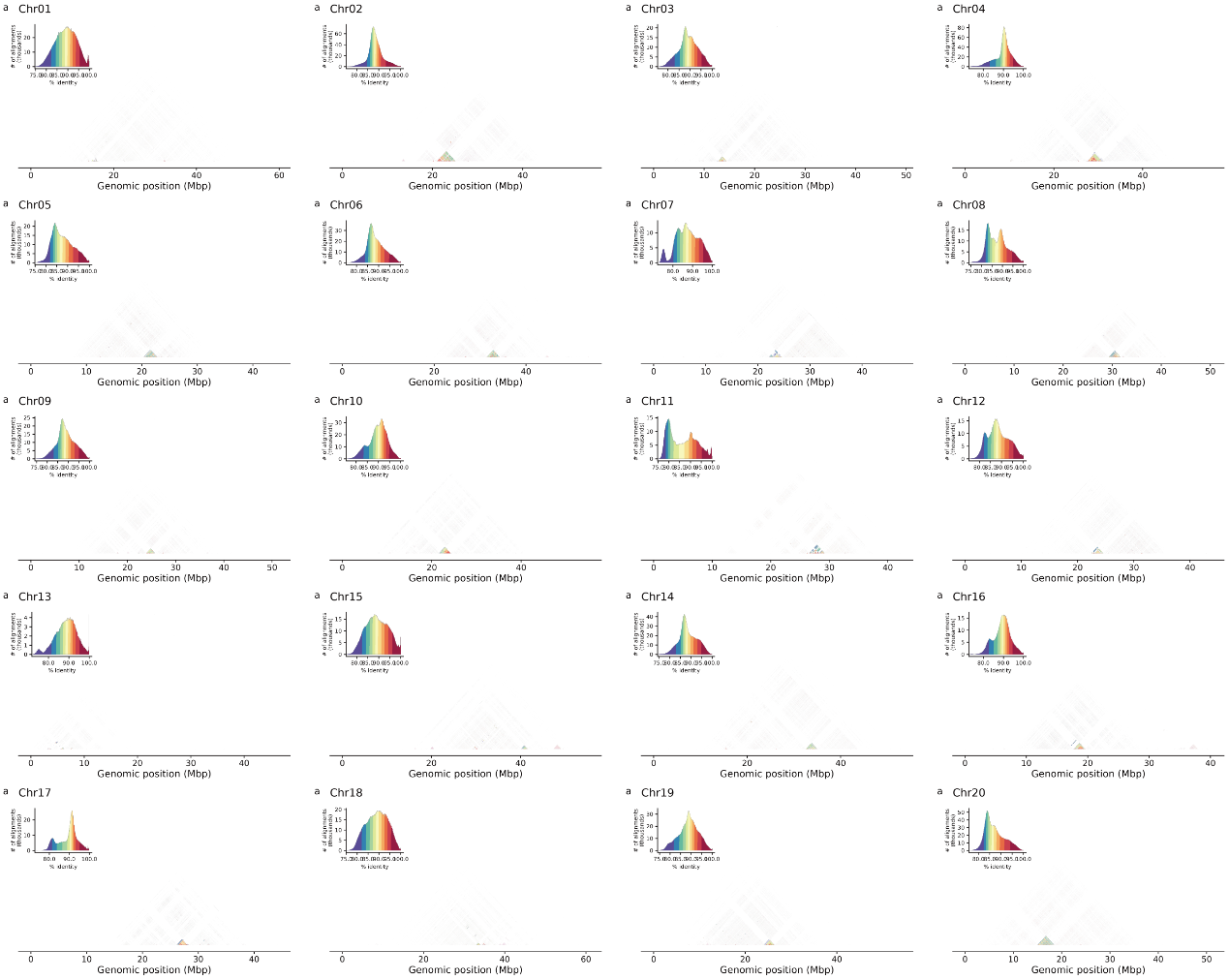

Fig. S14 Similarity of tandem arrays on soybean chromosomes, with clustered triangles representing TRAs and redder coloration indicating higher similarity.

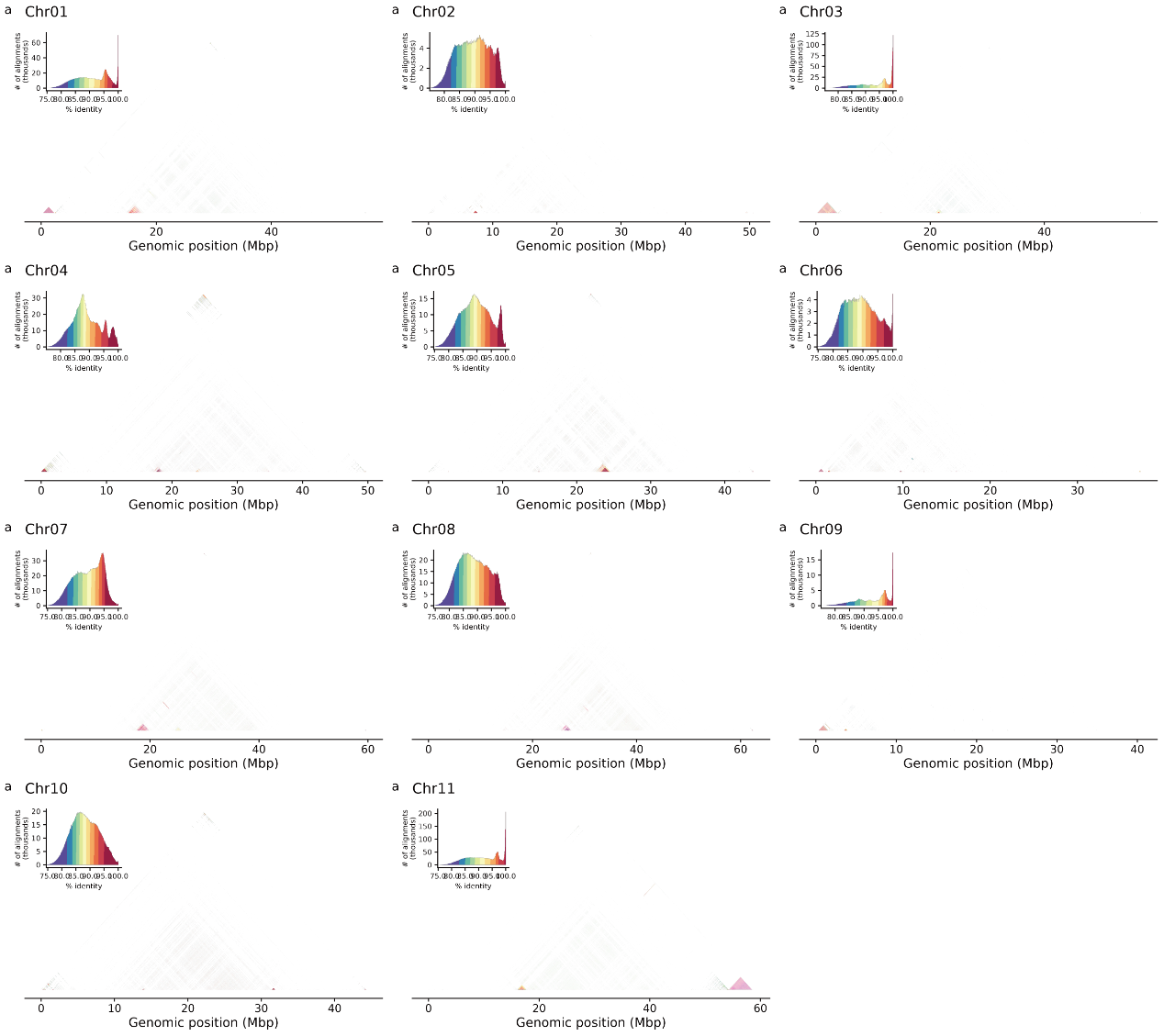

Fig. S15 Similarity of tandem arrays on common bean chromosomes, with clustered triangles representing TRAs and redder coloration indicating higher similarity.

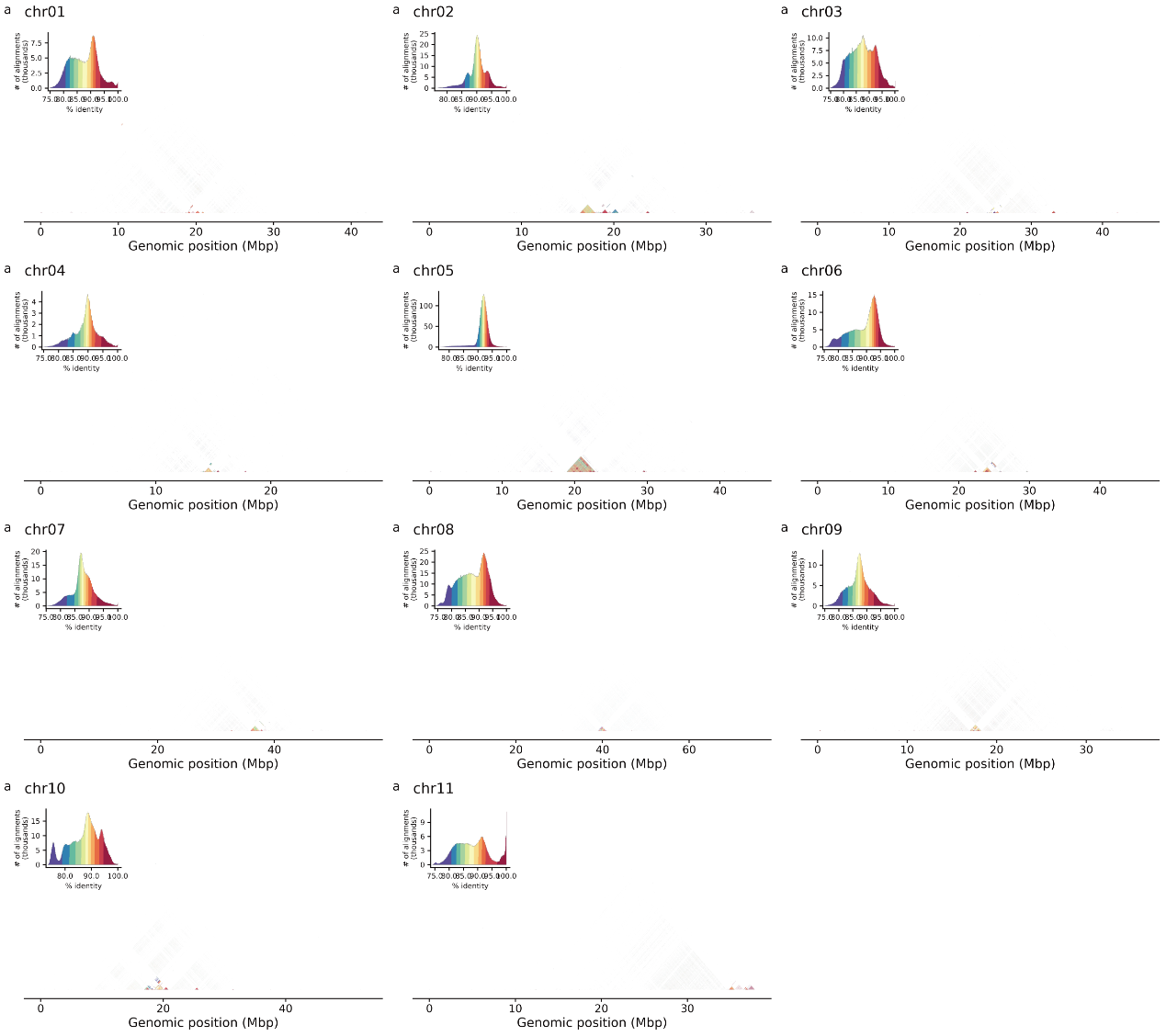

Fig. S16 Similarity of tandem arrays on mung bean chromosomes, with clustered triangles representing TRAs and redder coloration indicating higher similarity.

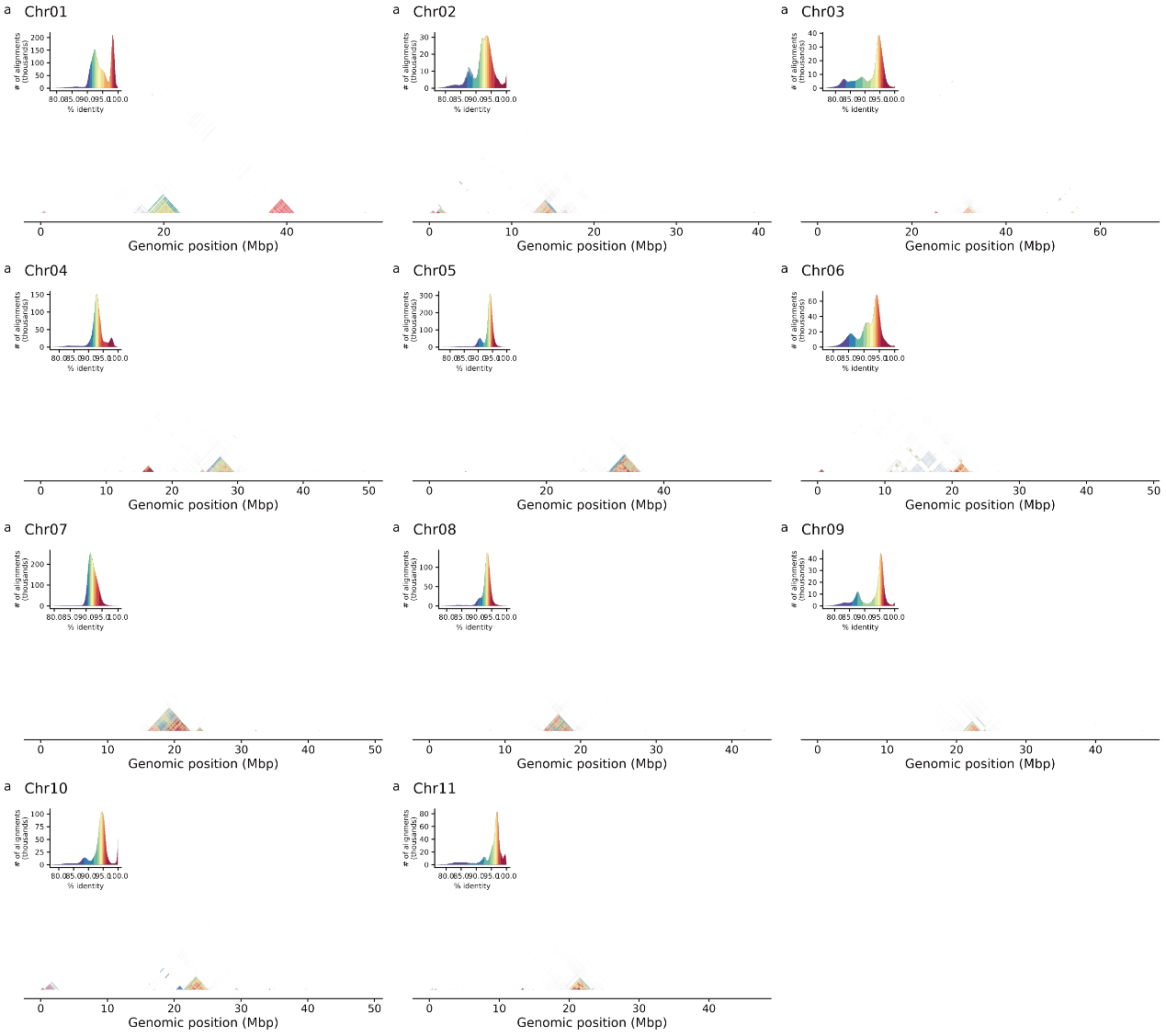

Fig. S17 Similarity of tandem arrays on cowpea chromosomes, with clustered triangles representing TRAs and redder coloration indicating higher similarity.

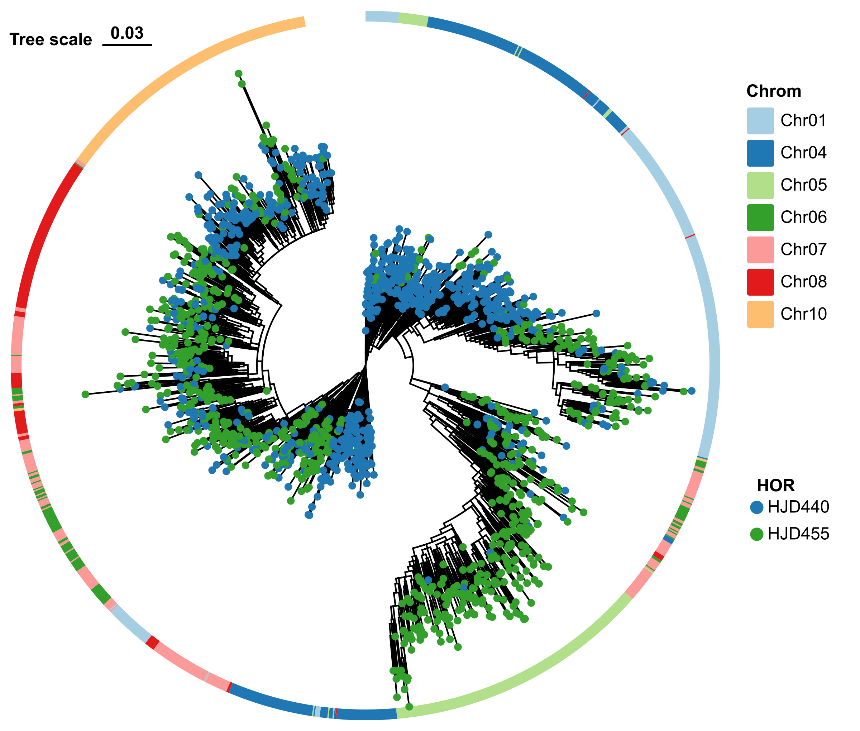

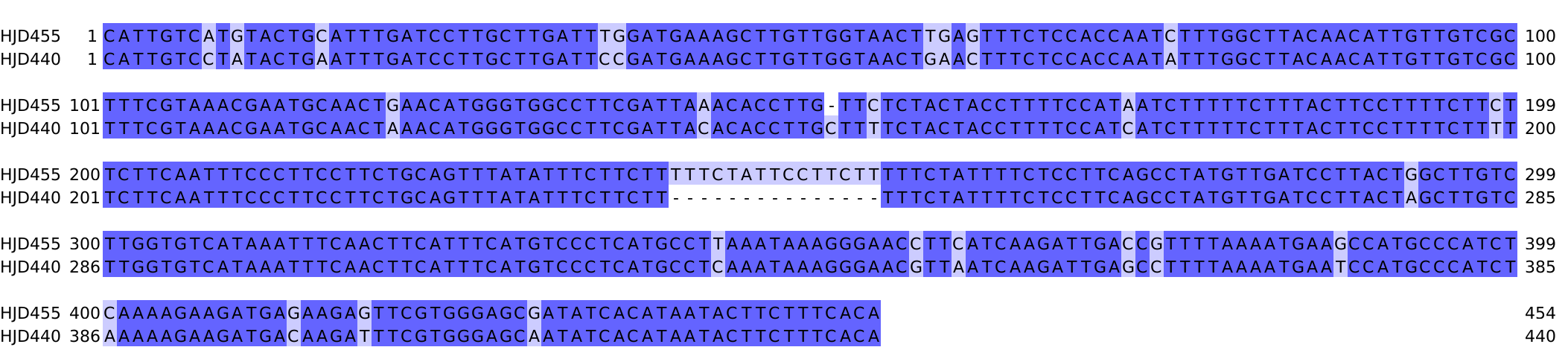

Fig. S18 Phylogenetic tree of selected CEN440 and CEN455 elements and corresponding sequence alignment of representative monomers.

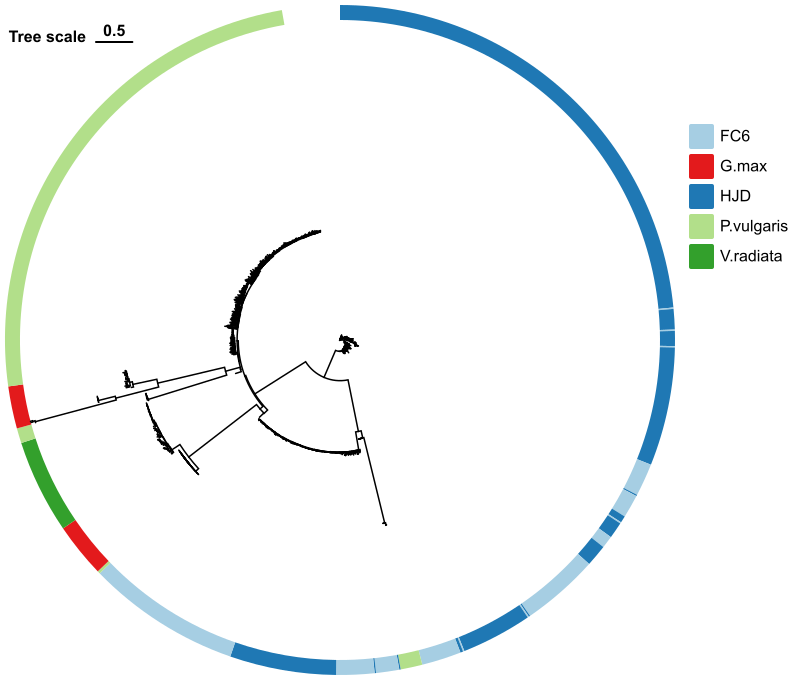

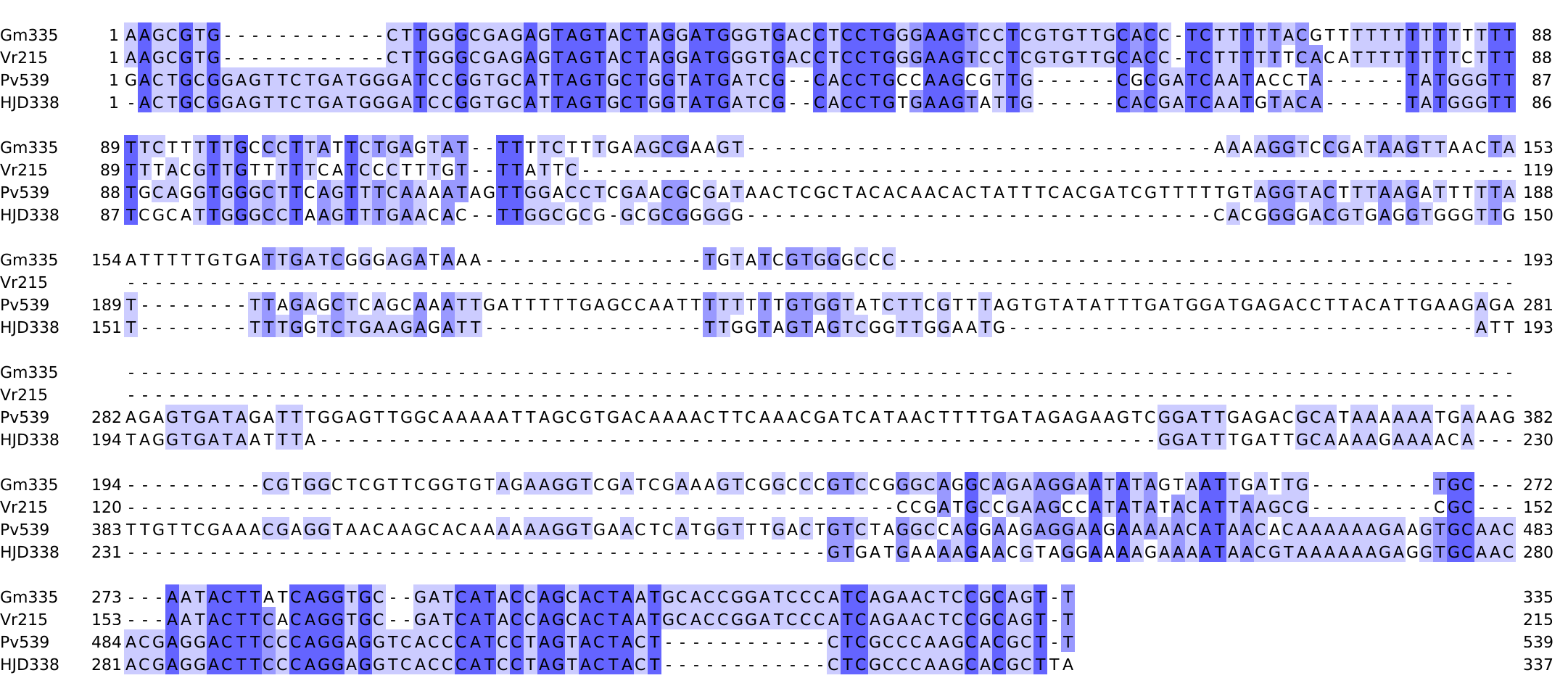

Fig. S19 Phylogenetic tree and representative sequence alignment of Gm335, Vr215, Pv539 and HJD338.

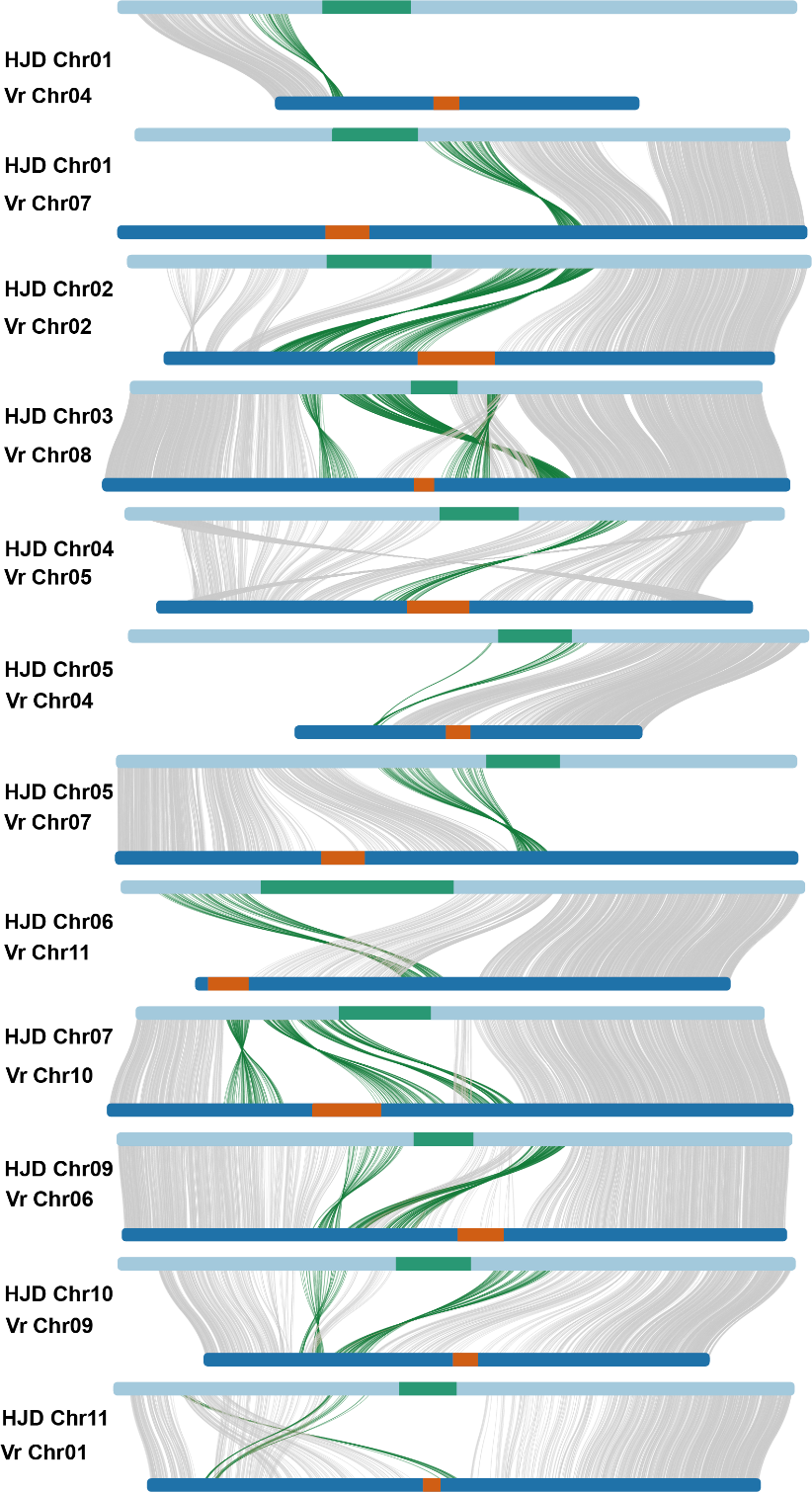

Fig. S20 Synteny between cowpea and mung bean; central lines connect homologous chromosomes, colored blocks mark centromeres, gray lines link homologous genes, and green lines highlight genes within or adjacent to centromeres.

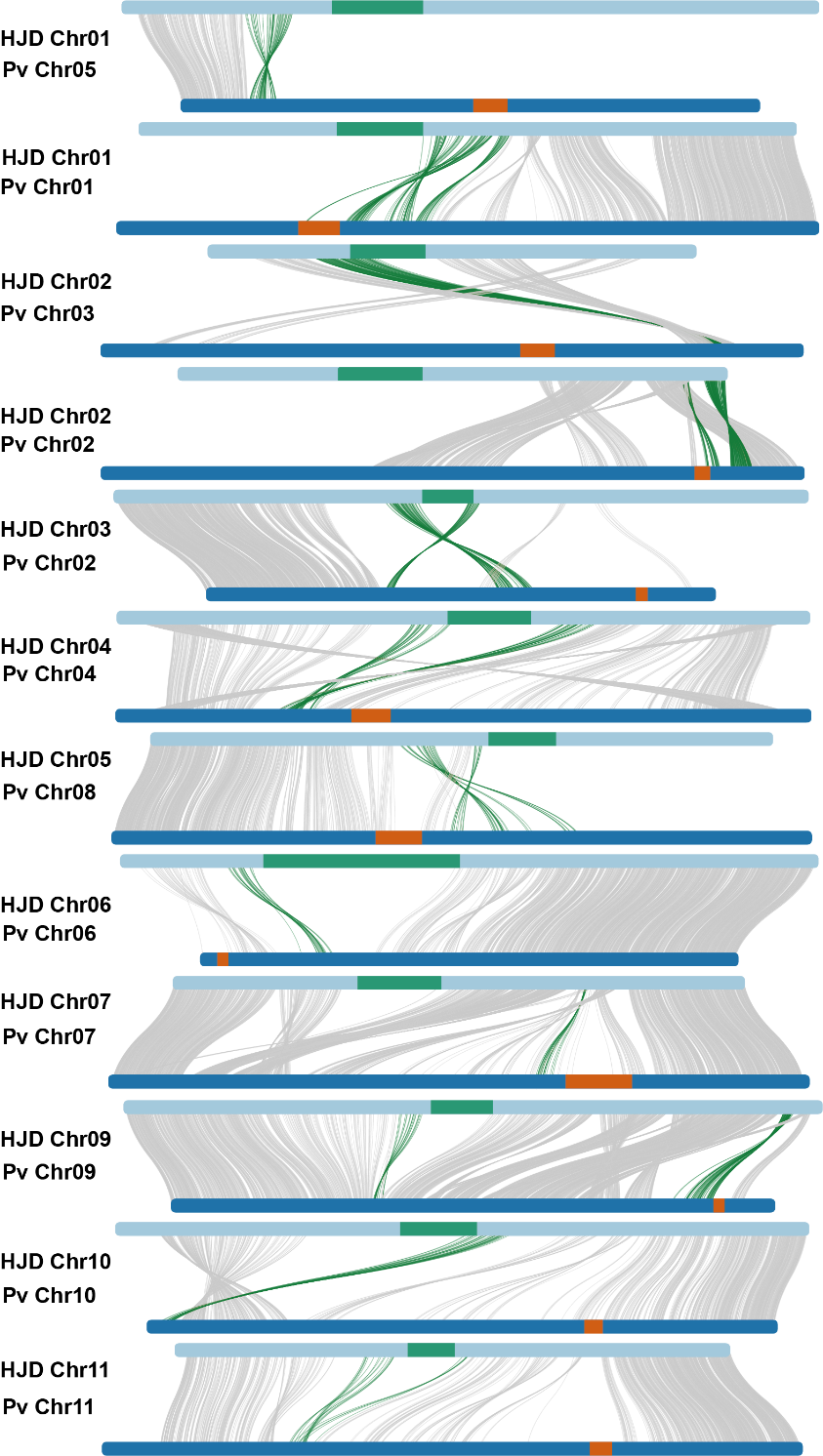

Fig. S21 Synteny between cowpea and common bean; central lines connect homologous chromosomes, colored blocks mark centromeres, gray lines link homologous genes, and green lines highlight genes within or adjacent to centromeres.

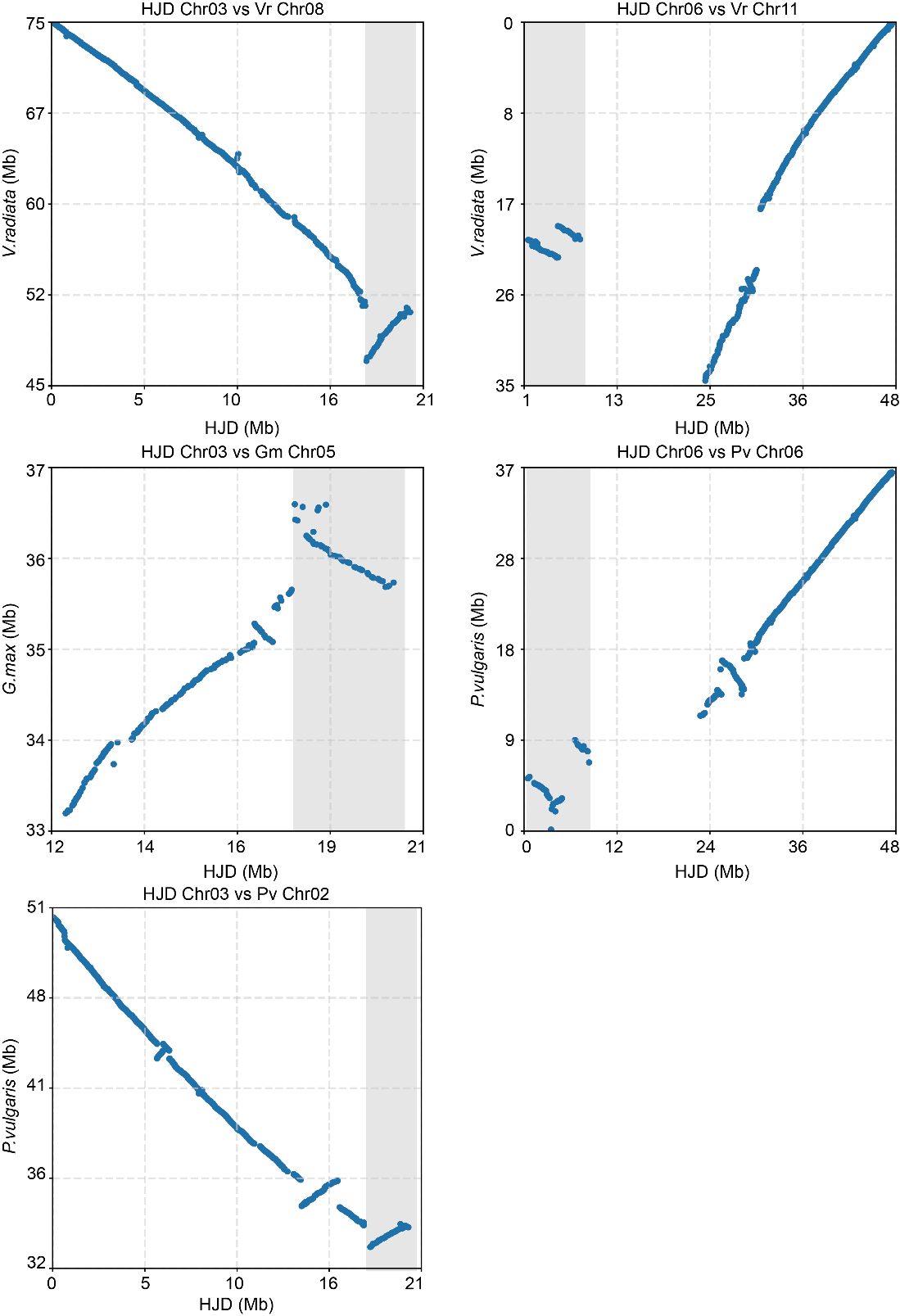

Fig. S22 Dot plots of inversions at shared breakpoints in cowpea versus Soybean, cowpea versus mung bean and cowpea versus common bean; blue dots represent orthologous gene pairs, gray shading indicates aligned regions.

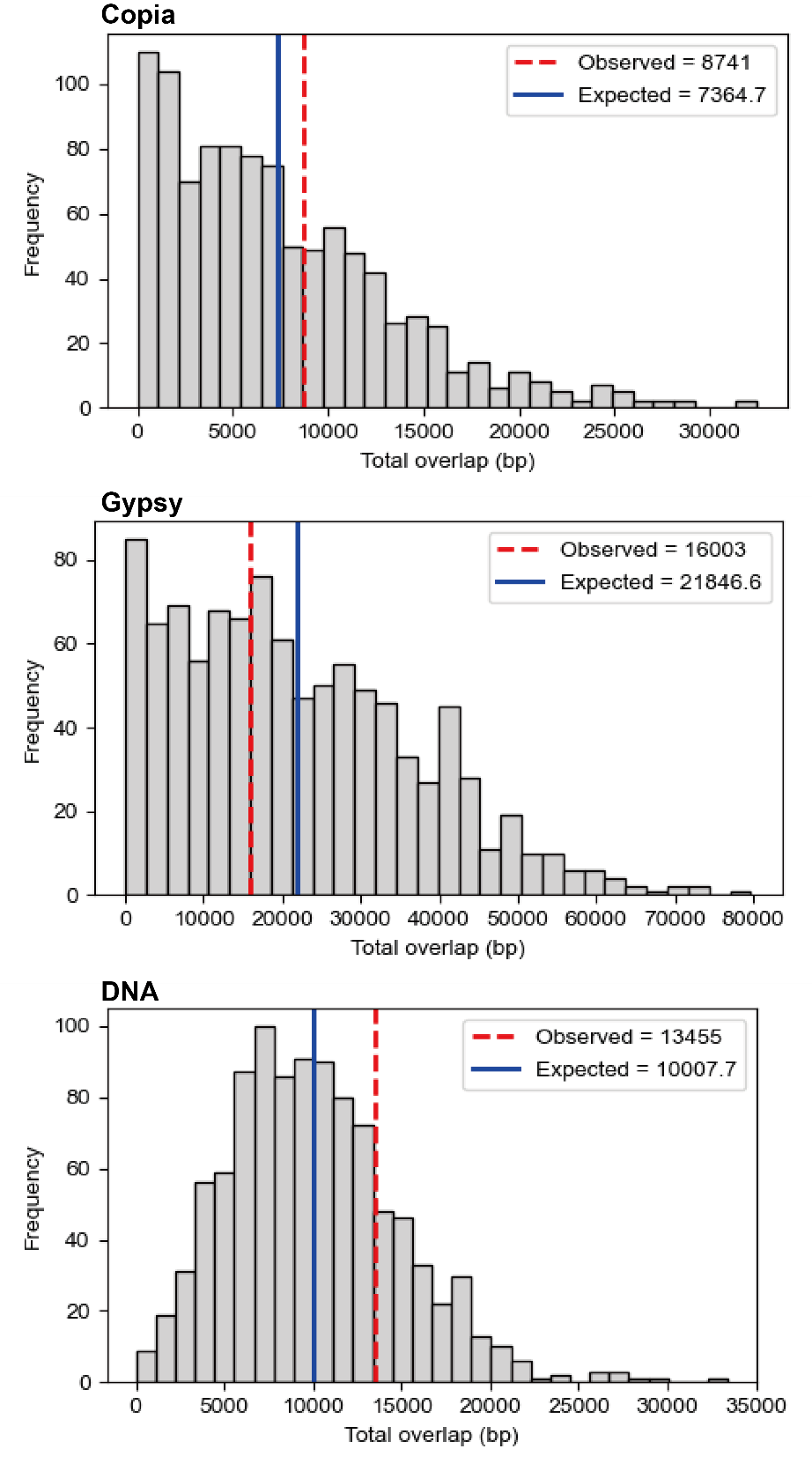

Fig. S23 Copia, Gypsy and DNA transposons occupancy at inversion breakpoints estimated from 1000 random resampling iterations.
